# Choice of climate data influences current and future global invasion risks for two *Phelsuma* geckos

**DOI:** 10.1101/2022.08.04.502765

**Authors:** Nicolas Dubos, Thomas W. Fieldsend, Markus A. Roesch, Stéphane Augros, Aurélien Besnard, Arthur Choeur, Ivan Ineich, Kenneth Krysko, Boris Leroy, Sparkle L. Malone, Jean-Michel Probst, Christopher Raxworthy, Angelica Crottini

## Abstract

Invasion risks may be influenced either negatively or positively by climate change, depending on the species. These can be predicted with species distribution models, but projections can be strongly affected by input environmental data (climate data source, Global Circulation Models and Shared Socio-economic Pathways SSP). We modelled the distribution of *Phelsuma grandis* and *P. laticauda*, two Malagasy reptiles that are spreading globally. We accounted for drivers of spread and establishment using socio-economic factors (e.g., distance from ports) and two climate data sources, i.e., Climatologies at High Resolution for the Earth’s and Land Surface Areas (CHELSA) and Worldclim. We further quantified the degree of agreement in invasion risk models that utilised CHELSA and Worldclim data for current and future conditions. Most areas identified as highly exposed to invasion risks were consistently identified (e.g. in Caribbean and Pacific Islands). However, projected risks differed locally. We also found notable differences in quantitative invasion risk (3% difference in suitability scores for *P. laticauda* and up to 14% for *P. grandis*) under current conditions. Despite both species native distributions overlap substantially, climate change will drive opposite responses on invasion risks by 2070 (decrease for *P. grandis*, increase for *P. laticauda*). Overall, projections of future invasion risks were the most affected by climate data source, followed by SSP. Our results highlight that assessments of current and future invasion risks are sensitive to the climate data source, especially in Islands. We stress the need to account for multiple climatologies when assessing invasion risks.

## Introduction

Invasive Alien Species (IAS) are increasingly raising concerns about their impact in the future, notably due to their rising economic cost and ecological impact (Diagne et al. 2021). Invasion risks can be influenced by climate change either positively or negatively depending on the species (Bellard et al. 2013). The assessment of invasion risks and how they will be influenced by climate change has become paramount to the development of proactive conservation actions.

Early detection is a key determinant to prevent invasions, suggesting the urgent need to identify priority areas for surveillance efforts. In this regard, a widely advocated management tool in conservation biology is Species Distribution Modelling (SDM; Gallien et al. 2012; Lanner et al. 2022). This approach consists of identifying the environmental factors explaining the distribution of a species and predicting areas of high environmental suitability. In the case of IAS that are expanding, such tools allow identification of areas that have not yet been invaded, but where environmental conditions are suitable and where factors of introduction and spread are present (e.g. maritime traffic). The distribution of IAS may be explained and predicted by a combination of environmental and socio-economic predictors (Bellard et al. 2016; Lanner et al. 2022). Environmental predictors (e.g. climate and habitat) may be used to identify areas where a candidate IAS is likely to further establish, while socio-economic predictors (e.g., proximity to ports and airports) represent factors of spread and entry points (Bertelsmeier and Courchamp 2014; Hulme 2021). The areas at high risk of invasion can then be prioritised for surveillance efforts.

Invasion risks may vary with climate change, and this can be assessed using SDMs that incorporate future climate projections (Bellard et al. 2013; Gillard et al. 2017). Such projections depend on the Shared Socio-economic Pathways (SSP, also known as Representative Concentration Pathway RCP, i.e., the scenario) and the Global Circulation Model (GCM, i.e., a methodological aspect). These factors can strongly affect SDMs and induce uncertainty in assessments of climate suitability (Buisson et al. 2010). More recently, the choice of the source of climate data (e.g., CHELSA or Worldclim; Fick and Hijmans 2017; Karger et al. 2017) used for model calibration has been identified as a major source of uncertainty in SDMs (Baker et al. 2016; Dubos et al. 2022a). In spite of this, most SDM studies aimed at projecting future distributions use climate data from a single source. To our knowledge, no study has considered the uncertainty induced by the source of climate data in invasion risk assessments.

Reptiles represent one of the costliest invasive taxa in terms of damage and management, with an estimated economic cost of more than one billion dollars per year in the last decades (Diagne et. al. 2021). The increasing pace of reptile invasions, along with the associated ecological (e.g., trophic disruptions), evolutionary/conservation (e.g., through hybridisation or introgression), and sanitary costs (e.g. pathogen transmission) have led to a growing attention towards some species (Reed and Kraus 2010; Sauteur et al. 2013; Kraus 2015; Vuillaume et al. 2015; Bellinati et al. 2022; Breuil et al. 2022). However, invasive alien reptiles remain understudied compared to invertebrate and plant species (e.g., Bellard et al. 2013) and there is a need to fill a knowledge gap in how climate change influences reptile invasion risks.

The Madagascar giant day gecko *Phelsuma grandis* Gray 1870 and the Gold-dust day gecko *Phelsuma laticauda* Boettger 1880 are two Malagasy reptiles that have spread throughout the world. *Phelsuma grandis* is one of the largest living species of the genus, reaching up to 30 cm in total length (i.e. twice the length of most *Phelsuma* species). *Phelsuma laticauda* is a medium-sized gecko, reaching up to 13 cm in total length; however, the species is considered an aggressive competitor towards other smaller gecko species (e.g., in French Polynesia, Lund 2015; but also in its native range, Gehring et al. 2010). Due to their human-mediated spread and resulting risks to native communities, both are considered IAS outside of their native range, including areas in central-eastern Madagascar, Mauritius, Reunion Island, Florida, French Polynesia and Hawaii (Ota and Ineich 2006; Krysko and Borgia 2012; Dubos 2013; Buckland et al. 2014; Dubos et al. 2014; Lund 2015; Fieldsend and Krysko 2019; Fieldsend et al. 2020, 2021c). The coexistence of *Phelsuma* spp. (or other reptiles sharing similar habitats such as anoles) may cause shifts in habitat use through competition (Harmon et al. 2007; Porcel et al. 2021; Wright et al. 2021) that might be detrimental to the more specialised native species. In Mauritius, the introduction *P. grandis* was associated with the extirpation of four populations of endemic *Phelsuma* species (Buckland et al. 2014). Both *P. grandis, P. laticauda* as well as several other *Phelsuma* species (e.g., *P. kochi*) are known to prey on other gecko specimens of smaller size (Gehring et al. 2010; Buckland et al. 2014; Rakotozafy 2019), which suggests potential predation risks to smaller species or juveniles of similar-sized species. Their introduction also raised concerns regarding the risk of disease and parasite transmission to native species (Dervin et al. 2014; Barnett et al. 2018; Fieldsend and Krysko 2019; Fieldsend et al. 2021b; Unger et al. 2022), despite no evidence of cross-species infection having been found so far (Goldberg and Bursey 2000; Leinwand et al. 2005). The spread of IAS can be facilitated by the international pet trade, as it is the case for *Phelsuma* spp. (Andreone et al. 2012; Masin et al. 2014; Stringham and Lockwood 2018; Pragatheesh et al. 2021). The spread of *P. grandis* and *P. laticauda* has led to increased attention regarding the conservation status of the native (and often endemic) fauna from Madagascar, Mauritius and Reunion Island (Dubos 2013; Buckland et al. 2014; Dubos et al. 2014). Their co-occurrence with native *Phelsuma* species has raised concerns regarding the long-term persistence of *P. lineata, P. serraticauda, P. inexpectata, P. borbonica, P. cepediana, P. guimbeaui, P. ornata*, and *P. rosagularis* (Andreone et al. 2003; Glaw and Vences 2007; D’Cruze et al. 2009; D’Cruze and Kumar 2011; Blumgart et al. 2017; Porcel et al. 2021). Both the IAS considered here are commonly found in urbanised areas, on ornamental plants and in orchards, as well as primary rainforests, reflecting a large niche flexibility that may help to explain successful establishments (Fieldsend et al. 2021a). This illustrates the need to characterise their climatic niche in order to identify potential areas at risk of invasion at the global scale.

Here we modelled the distribution of *P. grandis* and *P. laticauda* under current and projected future climatic conditions, and predicted their invasion risks at the global scale. To determine whether the choice of climate data source affects invasion risk assessments, we quantify the degree of agreement between current invasion risks based on the two main climate data sources available at global scale (i.e., CHELSA and Worldclim). We account for multiple sources of uncertainty for each climate data (Shared Socio-economic pathways, SSP and Global Circulation models, GCM). To provide the most reliable conservation guidelines, we identify areas that are in agreement between projections derived from both climate data sources, and point out priority areas for monitoring to enhance the chances of early detection and prevent potential invasions.

## Methods

Both *P. grandis* and *P. laticauda* are found in a variety of habitat types, including primary forests, highly degraded forests, orchards, and urbanised habitats (D’Cruze et al. 2009; D’Cruze and Kumar 2011; Dubos et al. 2014; Blumgart et al. 2017). We thus assume that habitat variables represent poor predictors of their environment, and that the species’ distribution may be better predicted by climate variables (e.g., Fieldsend et al. 2021a). Our analysis includes socio-economic factors such as proximity to roads, ports and airports, which we interpret as factors of spread or potential entry points.

### Occurrence data

We retrieved occurrence data from the literature and opportunistic observations, both from native and non-native ranges (Glaw and Vences 2007; Raxworthy et al. 2007; Pearson and Raxworthy 2009; Dubos 2013; Buckland et al. 2014; Dubos et al. 2014; Fieldsend and Krysko 2019; Fieldsend et al. 2021a; Fieldsend et al. 2021c; Porcel et al. 2021). In total, we obtained 338 unique occurrence records for *P. grandis* and 113 for *P. laticauda*. We thinned the data to avoid pseudo-replication and mitigate spatial biases, selecting one occurrence per pixel at the resolution of the environmental variables (5 arc minutes, see below). This resulted in a sample of 91 presence points for *P. grandis* based on CHELSA, of which 50 are within the native area and 41 in non-native areas (90 based on Worldclim; 49 and 41 points in native and non-native areas, respectively). For *P. laticauda*, the final sample represents 58 presence points, of which 19 are distributed in the native area and 39 in the non-native area (59 points based on Worldclim; 18 and 41 points in native and non-native areas, respectively).

### Climate data

We used 19 bioclimatic variables (description available at https://www.worldclim.org/data/bioclim.html) at 5 arc minutes (approximately 10 km) resolution for the current and future (2070) climate from two sources: CHELSA (Karger et al. 2017) and Worldclim global climate data (Fick and Hijmans 2017). These data sources used different methods to compute the climatologies. Worldclim is based on interpolated data with elevation and distance to the coast as predictors in addition to satellite data (Fick and Hijmans 2017), while CHELSA is based on statistical downscaling for temperature, and precipitation estimations incorporating orographic factors (i.e., wind fields, valley exposition, boundary layer height; Karger et al. 2017). We decided to include all 19 bioclimatic variables because both temperature and precipitation are related to the species’ biology, and use a statistical process to select the most relevant ones (see below). For each climate data source, we selected one predictor variable per group of inter-correlated variables to avoid collinearity (Pearson’s r > 0.7; Dormann et al. 2013) using the removeCollinearity function of the virtualspecies R package (Leroy et al. 2016). When mean values were collinear with extremes, we selected the variables representing extreme conditions (e.g., warmest / driest condition of a given period) because these are more likely to drive mortality and local extirpation, and be causally related to the species establishment (Parmesan et al. 2000; Mazzotti et al. 2016; Maxwell et al. 2019).

For future projections, we used three Global Circulation Models (GCMs; i.e., BCC-CSM1-1, MIROC5, and HadGEM2-AO) and two greenhouse gas emission scenarios (the most optimistic RCP26 and the most pessimistic RCP85) to consider a wide panel of possible invasion risk in 2070.

### Socio-economic factors

We used distance to port and airports as factors of introduction and proxies for propagule pressure (Bellard et al. 2016). We obtained port data from the World Port Index (https://msi.nga.mil/Publications/WPI, accessed December 2020) and airport data from the OpenFlights Airport database (https://openflights.org/data.html, accessed December 2020). We used distance to main roads and highways as an indicator of potential spread, since *Phelsuma* species can be accidentally transported by terrestrial vehicles over short distances (Deso 2001). We selected the largest two categories of road size (highways and primary roads) and computed distance from roads using the Global Roads Inventory Project (GRIP4) dataset (Meijer et al. 2018).

### Distribution modelling

We modelled and projected species distributions using an ensemble model approach (four modelling techniques). We selected a set of top-performing modelling techniques according to Valavi et al. (2021). These were Random Forest down-sampled (RF down-sampled, i.e., RF parametrised to deal with a large number of background samples and few presence records; Prasad et al. 2006), and three of the best performing models available in the biomod platform (Thuiller et al. 2009): a recent implementation of MaxEnt, i.e. MaxNet (Phillips 2017), Generalised Boosting Model (GBM, also known as Boosted Regression Tree, BRT; Elith et al. 2008) and Generalised Additive Model (GAM; Guisan et al. 2002). RF down-sampled was set to run 1000 bootstrap samples/trees.

Our dataset consisted of presence-only data. Hence, we generated pseudo-absences at locations where the species has never been detected (Sillero et al. 2021). We first generated five different sets of 50,000 randomly-selected pseudo-absences (or background points). Our occurrence data were retrieved from opportunistic observations, and were thus subject to spatial biases (e.g., more observations around populated or accessible areas). To account for sample bias, we reperformed all calculations applying a correction based on a different pseudo-absence generation strategy (both corrected and uncorrected models are needed to reliably measure the effect of sample bias correction; Dubos et al. 2021b; details below). In corrected models, we produced five sets of pseudo-absences concentrated around the presence points to reproduce the spatial bsi of the sample, following Phillips et al. (2009). We used a null geographic model (i.e., a map of the geographic distance to presence points) generated with the dismo R package (Hijmans 2012) and used it as a probability weight for pseudo-absence selection. This technique was deemed appropriate for IAS that are still expanding (i.e., not at equilibrium), because it reduces the generation of pseudo-absences in regions that are suitable but not yet invaded (e.g., Lanner et al. 2022). Since no independent data are available to assess the effect of sample bias correction, we used the Relative Overlap Index (ROI) based on Schoener’s D overlap (Dubos et al. 2021b). The ROI enables assessment of whether the effect of correction is negligible compared to the variability between model runs. It computes (1) the mean overlap between the uncorrected and the corrected predictions (i.e., the absolute effect of correction), and (2) the overlap between every pair of model replicates (between each pseudo-absence and cross validation runs, individually for each modelling technique, i.e., model stochasticity). We computed the ROI as follows:

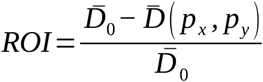

Where 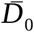 is the mean overlap between model runs of the corrected group and 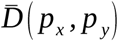 is the mean overlap between runs of the uncorrected and corrected models. A value close to 0 represents negligible effect of correction (i.e., the effect of sample bias correction is of same magnitude than model stochasticity). A value close to 1 represents a week effect of correction and strong model stochasticity. A negative value suggests that the correction effect is of lower magnitude than the model stochasticity and hence, irrelevant. We assumed that the correction affected our predictions if the overlaps between uncorrected and corrected groups were smaller than the overlaps between runs (i.e., ROI > 0; Dubos et al. 2021b).

We selected environmental predictors using a statistical approach, incorporating uncorrelated variables for which we had hypotheses of causality in the establishment or spread of the species. For each climate data source and species individually, we assessed the relative importance of each variable kept with 30 permutation per modelling technique (total = 120 per variable, data source, and species). The variables included in the final models were those with the highest relative importance. These were selected using the elbow criterion at the upper hinge of variable importance (i.e., the 25% best performing models per variable), setting a maximum of nine for *P. grandis* and five variables selected for *P. latiauda* following the ‘*number of observations m*/10 predictors’ rule-of-thumb proposed by Harrell et al. (1996) (see also Guisan and Zimmermann 2000). In total, we computed 400 models per species (4 modelling techniques × 5 pseudo-absence runs × 5 cross-validation runs × 2 modalities of sample bias correction × 2 climate data source) for the current distribution, and 2400 projections using future climate data (400 models × 3 GCMs × 2 SSPs).

### Model evaluation

Spatial partitioning is generally recommended to reduce spatial autocorrelation between training and testing data (i.e., block cross-validation; Valavi et al. 2019). In our case, occurrence data were highly aggregated, which results in strong unbalances between blocks. Therefore, we randomly partitioned the data, with 80% of the data being used for model calibration (training) and 20% for model evaluation (testing). This process was repeated five times (cross-validation runs) for each species, pseudo-absence dataset, correction modality, and climate data source. We assessed model performance using the Boyce index (Hirzel et al. 2006), assumed to be the best evaluation metric for pseudo-absence data (Leroy et al. 2018). A Boyce index value of 1 suggests that models predicted the presence points well, while a value of 0 means that model performance was not better than random. For ensemble models (i.e., the mean predictions across modelling techniques), pseudo-absence runs, and cross-validation runs for highly performing models, we discarded models for which the Boyce index was below 0.5.

### Quantifying the level of agreement in current invasion risks between climate data

Treating each species and sample bias correction modality separately, we compared the predicted current invasion risks obtained from CHELSA and Worldclim data. Firstly, we summed the suitability scores (total value of all pixels) of each ensemble model obtained and computed the absolute difference between CHELSA- and Worldclim-based predictions. This approach is the equivalent of the Species Range Change method (SRC; Buisson et al. 2010), except that we compared two projections of current distributions instead of two projections from different periods. The SRC indicates the overall level of agreement between two projections across the whole predicted area. A high SRC suggests a strong effect of the climate data source, either in terms of overall suitability scores or surface of suitable environment. We express the results as a percent absolute difference as follows:

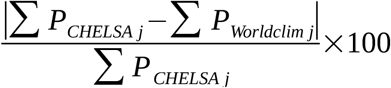

where *P*_*CHELSA*_ and *P*_*Worldclim*_ are the suitability score of pixel *j* for CHELSA and Worldclim projections, respectively.

Secondly, we used an approach that takes into account spatial information, i.e., spatial overlap (Muscatello et al. 2021; Petford and Alexander 2021; Dubos et al., 2022a). We computed the Schoener’s D overlap between projections of current invasion risk between predictions based on the two climate datasets considered. A value of 1 indicates a perfect spatial match between the two projections produced (i.e., no effect of climate data source) and a value of 0 represents a perfect mismatch. We computed the Schoener’s D overlap between CHELSA and Worldclim projections using the ENMTools R package (Warren et al. 2010). Schoener’s D was computed as follows:

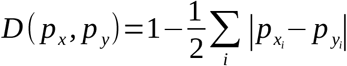

where *p*_*xi*_ and *p*_*yi*_ are the normalised suitability scores for uncorrected *x* and corrected *y* prediction in grid cell *i*, for each species, cross-validation run, and pseudo-absence run individually.

We quantified the uncertainty in SRC and Schoener’s D related to the climate scenarios, GCMs, and climate data source. For SRC (difference in summed scores), we quantified the proportion of deviance explained by climate data modalities using linear models (LM, assuming Gaussian errors), with SRC as the response variables, and the aforementioned sources of uncertainty as explanatory variables, following Baker et al. (2016). We then assessed the proportion of deviance explained by each source of uncertainty *f* as follows:

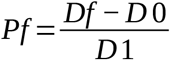

where, *Pf* = proportion of deviance explained by factor *f, D1* = deviance of full model, *Df* = deviance of full model minus factor *f*, and *D0* = deviance of null model.

We repeated this analysis for the Schoener’s D overlap using beta-regression GLM instead of LM, since overlap measures range continuously between 0 and 1 (glmmTMB R package; Brooks et al. 2019).

### Identifing priorities for surveillance

To identify areas at highest overall risk of invasion, we ranked countries and islands according to the invasion risk quantified for each species in the previous steps. We used border data obtained from the Global ADMinistrative area (GADM v4.0.4.; https://gadm.org/data.html) to associate the predicted invasion risks with the corresponding country, island, or archipelago (all territories with an ISO country code, e.g., Madagascar, Reunion Island, Comoros). We then computed the mean invasion risk per territory. To do so, we extracted the predicted values of our ensemble models and averaged them across all pixels of the territory. This approach may downplay the risks in large countries with only small regions at risk, but is useful for our study species which are mostly found in small islands. Since we had no a priori on which climate data source is best for ecological modelling, we based the ranking on the mean value between predictions obtained between CHELSA and Worldclim. To account for uncertainty, we penalised the mean prediction by subtracting its standard deviation (mean – SD), following the approach developed by Kujala et al. (2013) applied to single species (Dubos et al. 2022a). This enabled us to prioritise areas where the invasion risk is most-consistently identified as high across climate data sources and model replicates. For comparison, we also provide the rankings obtained from both individual climate data sources (tables available in supporting information).

### Projected effect of future climate change

We projected the predicted values of our models on future climate data. For each species, climate data, GCM, and scenario individually, we quantified the difference between current and future predicted distributions with the two aforementioned complementary approaches, i.e., difference in total (unpenalised) suitability scores and spatial overlap (Dubos et al. 2022a). We computed SRC as the difference between the summed suitability scores of future projections and current distribution and show the proportion of increase/decrease relative to current suitability scores, following Buisson et al. (2010) and Baker et al. (2016). Secondly, we quantified spatial suitability change using the Schoener’s D overlap to account for spatial information (for instance, allowing us to identify distributional shifts even when the total suitability does not change). We verified that models were well-informed for predictions on novel (future) data using clamping masks and examining the shape of predictor responses.

## Results

### Species distribution models

We selected six or seven variables for *P. grandis* depending on climate data source and five for *P. laticauda* (Table 1; Fig. S1–S8). The current distribution of both species was best explained by socio-economic variables ‘distance to ports’ and/or ‘distance to airports’ (Table 1) and climatic variables related to temperature variability (diurnal range bio2 and temperature seasonality bio4) or minimum temperature (bio6). Precipitation of warmest quarter (bio18) was also important for *P. grandis*. For this species, we discarded ‘distance to roads’ because this predictor produced spurious results, which did not correspond to our biological hypothesis (i.e., increasing invasion risk after 100 km distance) and would reduce the transferability of our models. Both species were present in the proximity of ports (approximately within 250km) and airports (approx. within 100km), in areas with low temperature variability, and with high minimum temperatures (> 10°C on average for coldest month) and in the case of *P. grandis*, avoiding dry regions (summer precipitation > 250mm; Fig. S3–S4, S7–S8).

**Table 1.**
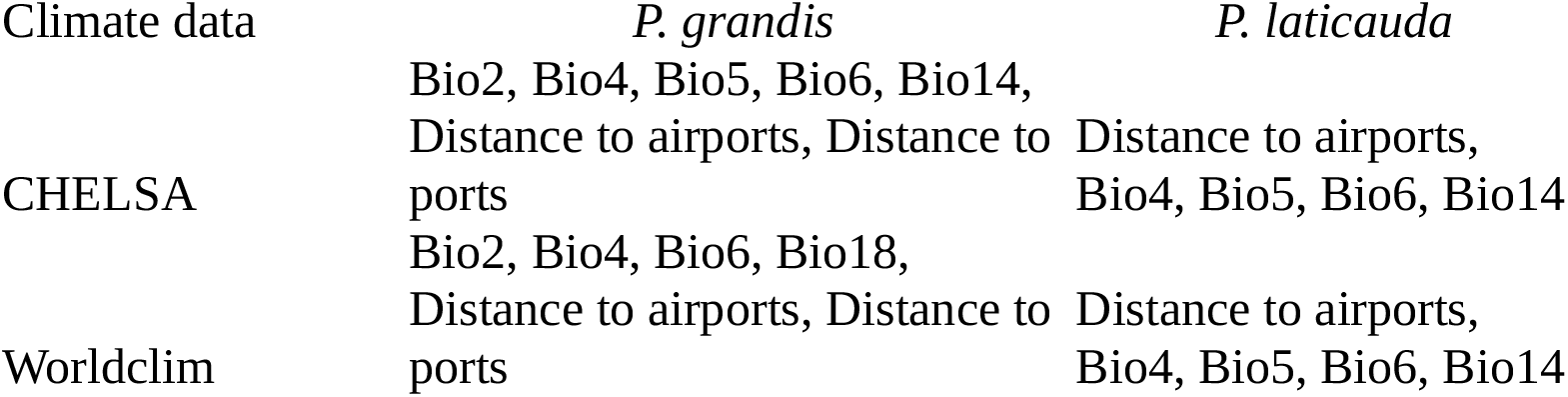
Selected environmental variables for the distribution modelling of two invasive *Phelsuma* species. Bio2: Temperature diurnal range; Bio4: Temperature seasonality; Bio5: maximal summer temperature; Bio6: minimal winter temperature; Bio14: precipitation of driest month; Bio18: summer precipitation.

Models generally did a good job of predicting known presences (most Boyce indices > 0.5), with higher Boyce indices for Worldclim-based predictions and corrected models compared to CHELSA-based and uncorrected models, respectively (Table 2; Fig. S9–S10). The effect of sampling bias correction was more important than model stochasticity (ROI = 0.07 with CHELSA, ROI = 0.09 with Worldclim for both species). We discarded 5 to 25 poorly performing models out of 100 per modality (species, climate data source, and sample bias correction; Table 2)

**Table 2.**
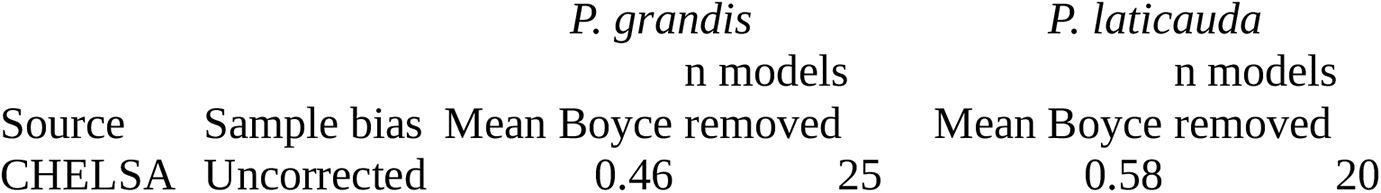
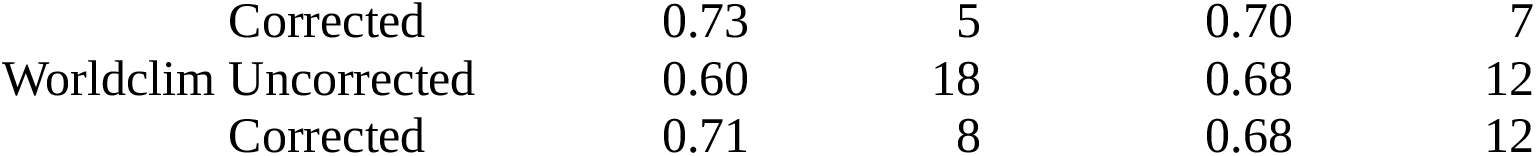
Model performance of the distribution models for two invasive *Phelsuma* species, inferred from mean Boyce indices. We show the number of poorly performing models (Boyce < 0.5) which were removed from the ensemble models (out of 100 models per modality).

|  |  | *P. grandis* |  | *P. laticauda* |  |
| --- | --- | --- | --- | --- | --- |
| Source | Sample bias | Mean Boyce | n models removed | Mean Boyce | n models removed |
| CHELSA | Uncorrected | 0.46 | 25 | 0.58 | 20 |
|  | Corrected | 0.73 | 5 | 0.70 | 7 |
| Worldclim | Uncorrected | 0.60 | 18 | 0.68 | 12 |
|  | Corrected | 0.71 | 8 | 0.68 | 12 |

### Current invasion risks

We identified important invasion risks in multiple regions throughout the world, mostly in tropical islands (Fig. 1, 2). In both species, we found the highest invasion risks in islands of the Indian Ocean (e.g., Comoros, Mayotte), Pacific Ocean (e.g., Niue, New Caledonia), the Caribbean region (especially in the Greater Antilles and the Bahamas), both coasts of central Africa (Angola, Congo, Tanzania, Mozambique), and the Indo-Pacific region (Philippines, Vietnam, New Guinea). Suitable conditions are also met in Cape Verde and the coast of Brazil for both species. We found different invasion risks between both species in the Lesser Antilles, Vanuatu, and the Hawaiian archipelago, with greater invasion risks for *P. laticauda* in these regions (Fig. 3).

**Fig. 1.**
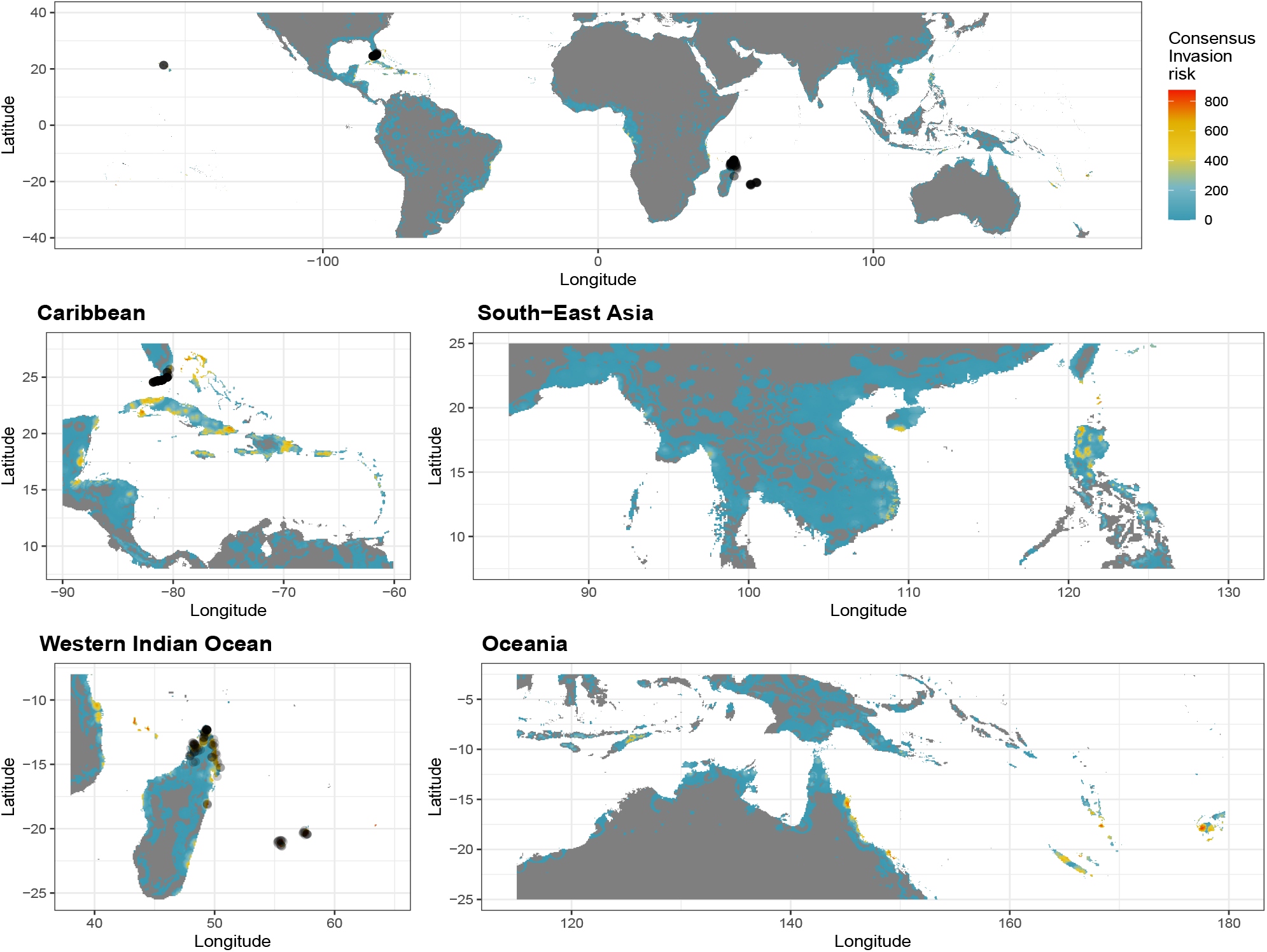
Current consensus global invasion risks for *Phelsuma grandis*. Consensus invasion risk was obtained from mean predictions between all simulations (including models based on CHELSA and Worldclim climate data), subtracting the standard deviation to account for uncertainty. Closed black circles represent the presence points. Climate data source-specific maps are available in supporting information (Fig. S11–S14).

**Fig. 2.**
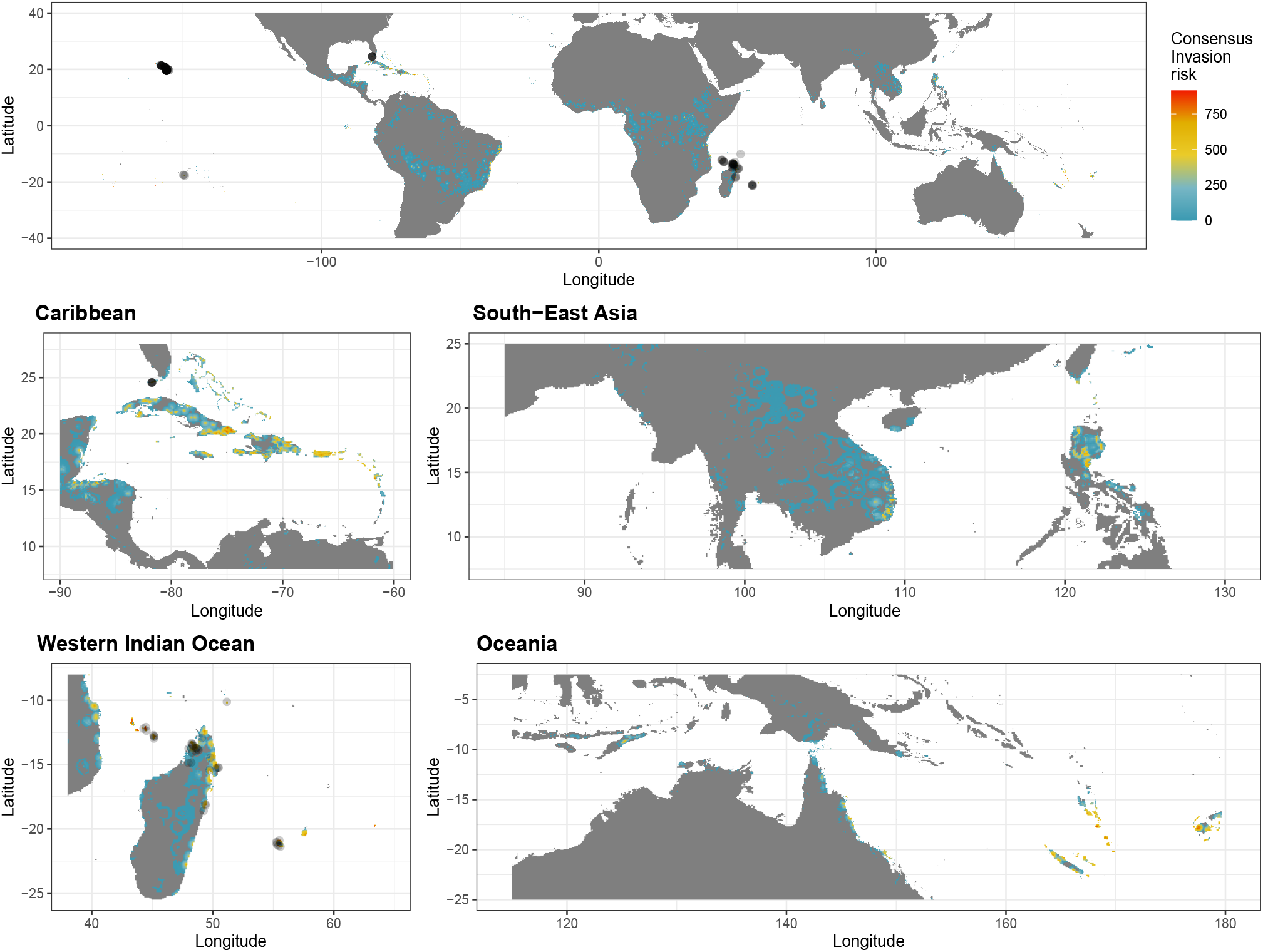
Current consensus global invasion risks for *Phelsuma laticauda*. Consensus invasion risk was obtained from mean predictions between all simulations (including models based on CHELSA and Worldclim climate data), subtracting the standard deviation to account for uncertainty. Closed black circles represent the presence points. Climate data source-specific maps are available in supporting information (Fig. S15–S18).

**Fig. 3.**
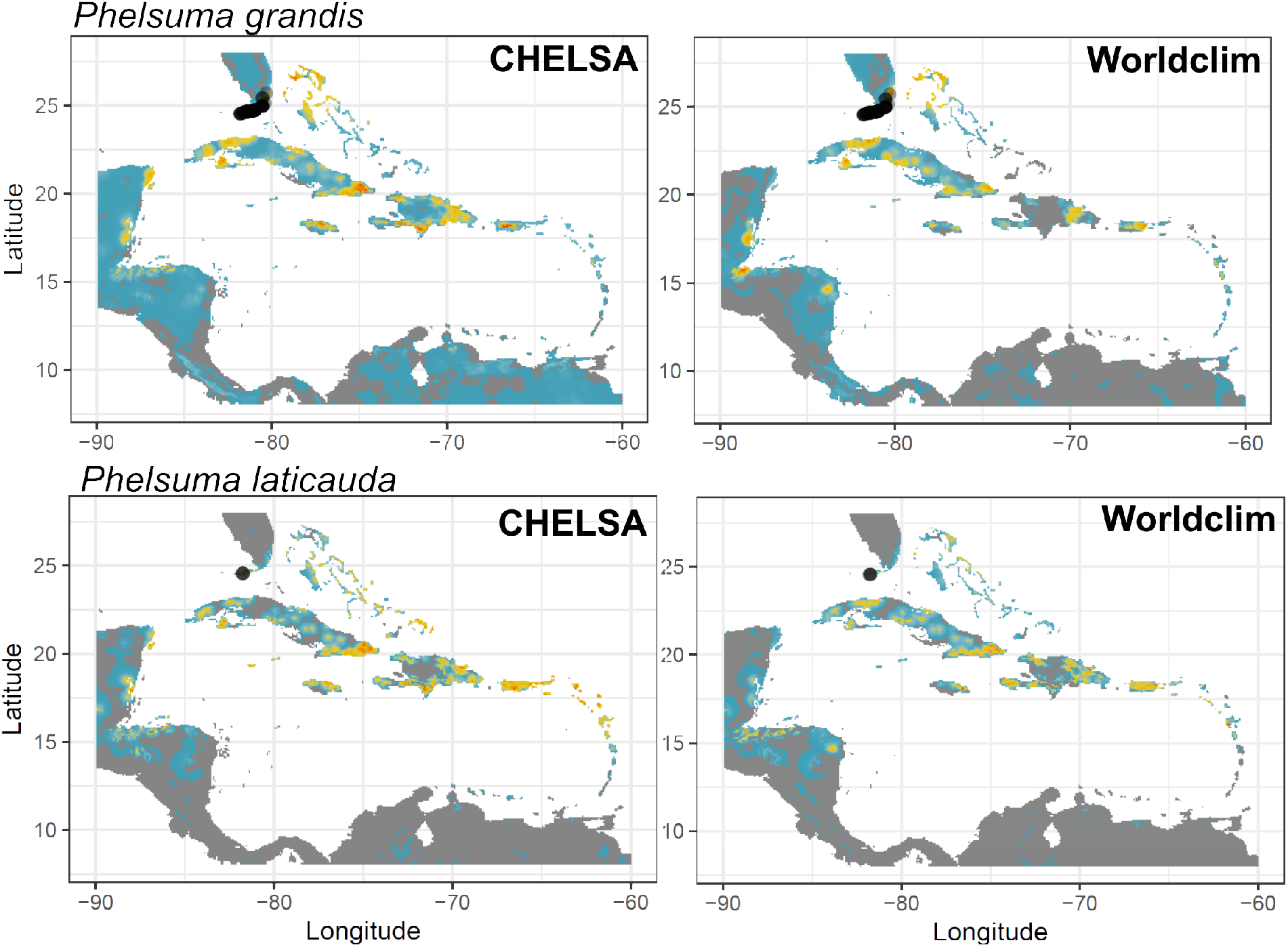
Predicted invasion risks of *Phelsuma grandis* (top) and *Phelsuma laticauda* (bottom) based on two sources of climate data (left: CHELSA; right: Worldclim) in the Caribbean region. Predicted values are averaged across model replicates (n = 100 per species and climate data source) and are penalised by uncertainty (standard variation across replicates). Red represents high invasion risk with high certainty, blue represents moderate invasion risk and grey represents areas where uncertainty was higher than invasion risks.

Projections of current invasion risks differed locally when calibrated using the different climate sources (see example of the Caribbean in Fig. 3; individual ensemble models are available in supporting information, Fig. S11–S18). We found notable differences between both projections in summed suitability scores and spatial overlap (Table 3). Climate data source affected the ranking of invasion risk per territory, with sometimes dramatic differences (e.g., 36 rank difference for Saint Helena, Ascension, and Tristan da Cunha for *P. grandis*; 30 rank difference for Saint Martin for *P. laticauda*; Table S1, S2).

**Table 3.** Degree of agreement between current invasion risks assessed from CHELSA and Worldclim climate data. We computed the absolute difference between the summed suitability scores (expressed as percentage), and Schoener’s D spatial overlap (expressed as spatial difference: 1-D). A higher value suggests a lower agreement.

|  | Difference in<br>scores (%) | 1-Schoener's<br>D overlap<br>(%) |
| --- | --- | --- |
| *P. grandis* | 13.9 | 17.5 |
| *P. laticauda* | 2.8 | 12.0 |

### Future invasion risks

We predict a decrease in invasion risk by 2070 in most cases (Fig. 4; S19–22). We found important differences between projections based on different climate data, with higher SRC for Worldclim-based projections overall (Fig. 4). For *P. grandis*, we predict a decrease in total invasion risk in all cases, ranging between 8.6 and 16.1 % (total scores), and a spatial change ranging between 9.9 and 19.4 % (1-Schoener’s D overlap) depending on the scenario and the climate data. For *P. laticauda*, the effect of climate change differed between climate data sources, ranging between -4.7 % (decline) and +18.8 % (increase) in overall invasion risk, and a spatial change ranging between 11.7 and 20.8 %. Clamping masks indicated novel conditions for one variable throughout the native and invaded range, for Worldclim only (Fig. S23, S24). These novel conditions seem to be mostly driven by maximal temperatures (Bio5; Fig. S25). Given the shape of the relationships between this variable and suitability (Fig. S3, S7, S8), there is little risk of uncertain predictions due to extrapolation.

**Fig. 4.**
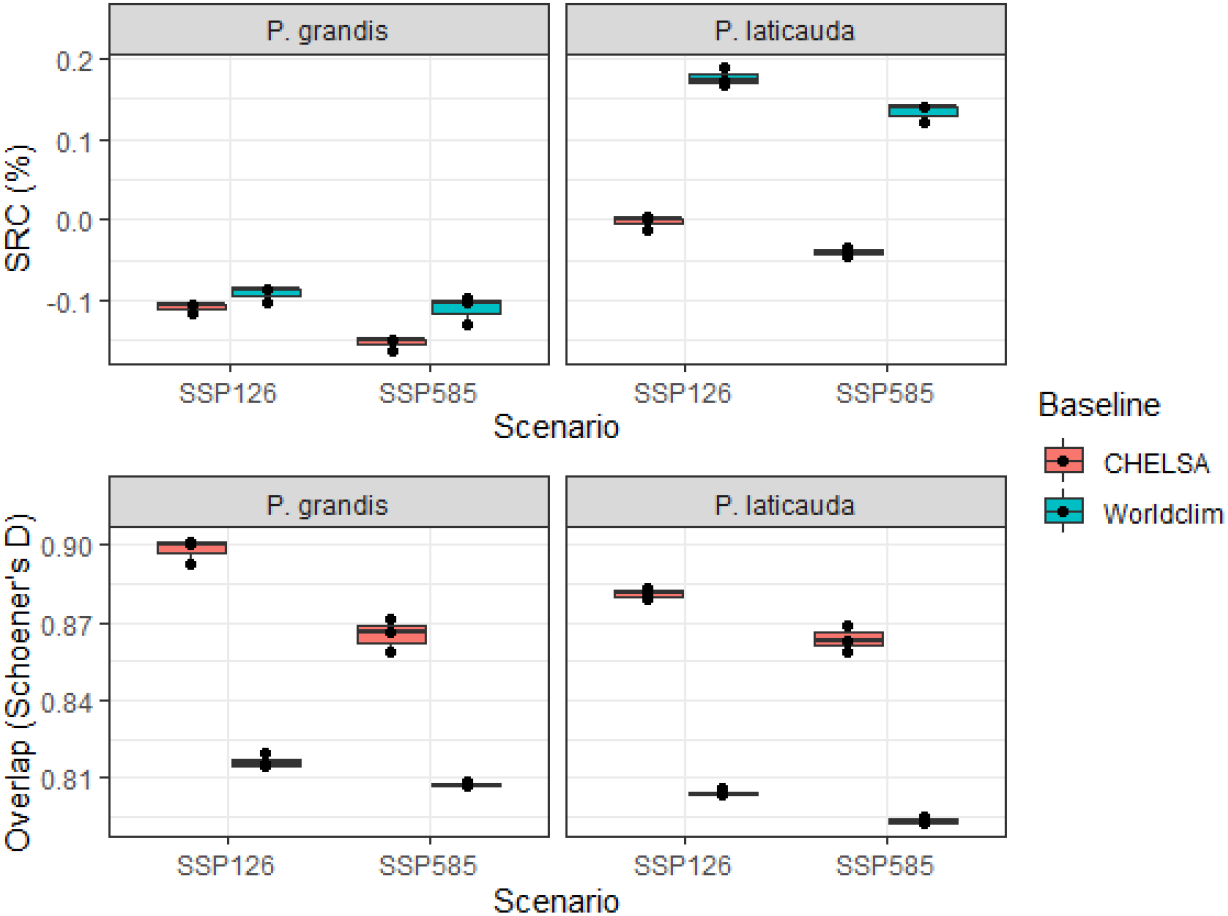
Effect of climate data source on Species Range Change (SRC, overall difference between current and future suitability, here expressed as a proportion relative to current suitability) and Schoener’s D overlap (percentage of common spatial information between current and future projections) for the invasive *P. grandis* and *P. laticauda*. Models were corrected for sample bias. Indices were computed individually for each climate data source, scenario (SSP) and Global Circulation models (GCMs, represented by the black points). Boxes represent the 25^th^ and 75^th^ percentile and the bars represent the median.

In most cases, the source of climate data was the most important driver of uncertainty in future invasion risk projections (Fig. 5). In terms of total scores (SRC), the source of climate data explained as much uncertainty as the scenario for *P. laticauda*. Otherwise, climate data source was by far the largest driver of uncertainty in terms of spatial overlap for both species.

**Fig. 5.**
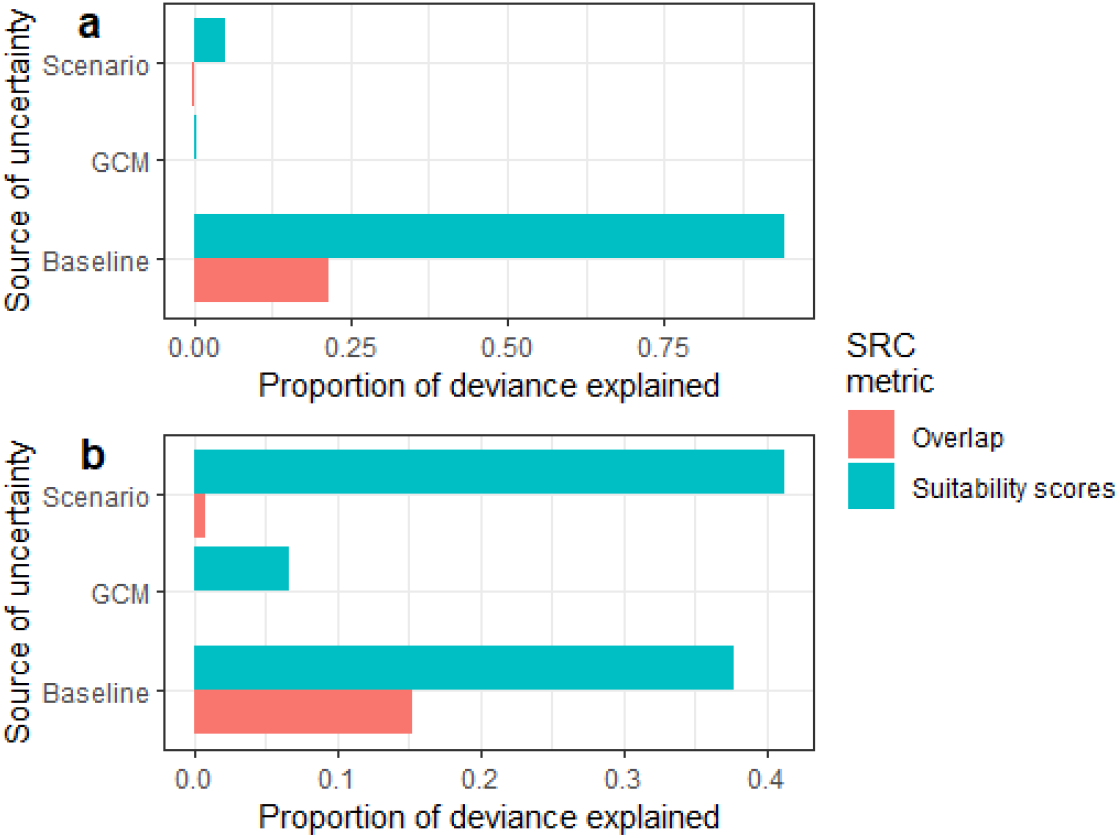
Proportion of deviance explained by climate data modalities (Scenario, Global Circulation Model and climate data source) in climate change effect (Overlap and Suitability scores) on (a) *Phelsuma grandis* and (b) *Phelsuma laticauda*. Overlap was computed with Schoener’s D between current and future projections; Suitability scores are the difference in total scores between current and future projections.

## Discussion

We modelled the current and future distribution of two invasive alien reptile species using a recently advocated approach, accounting for socio-economic factors and a wide panel of climate data. We identified several areas at high risk of invasion, findings that were robust to the choice of climate data. We propose that these areas be considered priority areas for surveillance efforts and monitoring, but areas identified at risk by single climate data must be also considered. We found important differences relating to the source of climate data. Overall, climate change will reduce invasion risks for *P. grandis* and slightly increase for *P. laticauda*.

### Drivers of spread and establishment

The spread of both species is driven by maritime and/or aerial transportation. The importance of proximity to ports and airports may be caused by the insularity of our study species (ports and airports are present in most islands). However, with our pseudo-absence sampling strategy (being more concentrated around presence points), the detection of this effect of these variables suggests that species are more present near ports and airports than random. Both *P. grandis* and *P. laticauda* are commonly found on anthropogenic structures including hotels and plant nurseries (pers. obs., Ineich, Choeur, Crottini; Gehring et al. 2010), as well as ornamental plants and plantation crops such as bananas and coconuts (Gill et al. 2001; D’Cruze et al. 2009; Porcel et al. 2021). Hence, individuals may be regularly carried via containers and accidentally introduced into shipments (Fritts 1987; Dubos et al. 2014; Khoury et al. 2021). Introductions were also caused by intentional or accidental release from captivity in regions where they are imported for the pet trade (e.g., in Florida; Andreone et al. 2012; Fieldsend and Krysko 2019).

Both species were able to establish in areas with a low variability in temperature, both at the daily and annual scale. Their native range is located at the north of Madagascar, close to the shores, where the climate is mainly equatorial (Peel et al. 2007), with little annual and daily thermal variation. Their affinity with low thermal variability may be related to the strong effects of temperature fluctuation on activity and reproduction (Georges 2013; Noble et al. 2018. Choeur et al. 2022). Our study species are active throughout the year, which may explain their affinity with low temperature seasonality. Their year-round activity implies a need for continuous food availability. A low seasonality may serve to maintain fruit/nectar production and insect activity, all sources of food for both *P. grandis* and *P. laticauda* (Dervin et al. 2013; Dubos et al. 2020; Hoarau et al. 2021). The link with low variability at the daily and annual scale may be due to temperature-dependant sex determination, a common feature in reptiles. Both, *P. grandis and P. laticauda* lay their eggs on the surface of the substrate, exposing them to daily fluctuation in temperature. In our case, a relatively constant temperature during the day and throughout the year may help balancing the sex-ratio and maintain population dynamics (Georges 2013).

We found that our study species were not able to establish in regions with low minimal temperature (< 10°C), presumably because the cold reduces the activity of ectotherms and hence, their survivability. This corresponds to the lower bound of thermal tolerance commonly found in tropical reptiles (Sunday et al. 2011). Minimal temperature may also influence incubation duration and sex determination (Georges 2013; Roesch et al. 2021). An extended incubation period may increase the probability of hatching failure and egg predation. Lower nesting success and unbalanced sex ratio could disrupt population dynamics and prevent persistence in colder regions. The establishment of both species in Florida may be surprising given the low temperatures occasionally occurring during winter compared to northern Madagascar. Recent assessments of the climatic niche of *P. grandis* revealed an important dissimilarity between the climate of its native range and the invaded areas of Florida (Fieldsend et al. 2021a). This suggests a high potential for either thermal plasticity or adaptation to new environments (Card et al. 2018; Lapwong et al. 2021), or underlies that the species’ native distribution is strongly limited by biotic interactions (predation and competition; e.g., competition with *P. kochi*; Fieldsend et al. 2021a). This is consistent with findings on *Hemidactylus frenatus* and *Anolis sagrei* which were able to spread in areas colder than their native range (Angetter et al. 2011; Lapwong et al. 2021). Invasive success is often facilitated by high genetic diversity (Angetter et al. 2011), which may be enhanced by multiple native-range sources as it is the case for *P. grandis* in Florida (Fieldsend et al. 2021c). Further research may assess the level of genetic diversity of both species throughout their invaded range to better understand the species ability to persist in new environments.

Both gecko species did not establish in regions with arid seasons, presumably because low precipitation limits primary and secondary production and therefore food availability (Dubos et al. 2019). Prolonged drought periods are associated with body condition declines, increased mortality, and local extirpation in reptiles (Maxwell et al. 2019), conditions which may prevent the establishment of our study species.

### Current invasion risks

Areas predicted to be at high risk of invasion were consistent between CHELSA- and Worldclim-based projections, but with locally important differences. These were mostly located in islands of the Caribbean, the islands of the Western Indian Ocean, South-East Asia, and Eastern Oceania. The potential establishment of invasive alien *Phelsuma* species in these areas may expose the local fauna to new competitors or predators. Both *P. grandis* and *P. laticauda* are highly flexible in terms of habitat use (D’Cruze and Kumar 2011; Dubos et al. 2014), which raises concerns for native synanthropic species as well as for species dwelling in natural forested habitats. Species at risk include native *Phelsuma* species (as suggested by the reduction of the *P. lineata* population in the eastern seaport town of Toamasina), or any other diurnal arboreal reptiles with similar habitat use (e.g., perch height, substrate; Augros et al. 2018; Wright et al. 2021), such as the Critically Endangered brown red-bellied anole *Anolis koopmani* from Haiti, the Endangered black-throated stout anole *Anolis armouri* from Haiti and the Dominican Republic, or the Critically Endangered Finca Ceres anole *Anolis juangundlachi* from Cuba. Given the broad range of habitat types occupied by our study species, conservation concern should also be given to all smaller species for which distribution matches the areas at risk (e.g., *Bavaya* spp. or *Eurodactylodes* spp. from New Caledonia). The Critically Endangered ‘Eua Forest Gecko *Lepidodactylus euaensis* from Tonga is of particular concern, given its conservation status and the very high invasion risk identified for this island. Both *P. grandis* and *P. laticauda* are diurnal, but can also be active at night due to artificial light (Dubos et al. 2020; Baxter-Gilbert et al. 2021), highlighting the risk of competition with nocturnal species living near anthropogenic structures such as the Critically Endangered Barbados leaf-toed gecko *Phyllodactylus pulcher* (Williams et al. 2016). The potential impact of invasive *Phelsuma* species on native fauna may be mitigated by potential plasticity, which could promote microclimatic and/or habitat partitioning (Noble et al. 2011; Porcel et al. 2021; Ryan and Gunderson 2021). Future studies should investigate the potential for spatial, temporal, or environmental shift for *P. grandis, P. laticauda*, and their sympatric species to better understand which species are at greater risk.

### The choice of climate data source

We identified local differences between predicted invasion risks using different climate data sources. Differences may be driven by the selection of different variables (e.g., models calibrated with CHELSA data selected ‘Daily temperature range’ but not with Worldclim for *P. grandis*); however, a recent study showed that differences can persist even when the same predictors are selected (Dubos et al. 2022a; see also Jiménez-Valverde et al. 2021). The mismatch may be better explained by the methods used to compute the climatologies. Worldclim is built from interpolated data with elevation and distance to the coast as predictors in addition to satellite data (Fick and Hijmans 2017). CHELSA used statistical downscaling for temperature, and precipitation estimations incorporate orographic factors (i.e., wind fields, valley exposition, boundary layer height; Karger et al. 2017). Such differences may be exacerbated in areas with strong topographic heterogeneity such as Oceanic Islands (e.g., Lannuzel et al. 2021). The difference in temporal coverage may represent another source of mismatch, with Worldclim representing the conditions of the 1960–1990 period while CHELSA was computed for 1979–2013. Since we have no a priori knowledge of which climate data source is most useful for predicting invasion risks, we suggest that studies aiming to predict current and future invasion risks should consider multiple climate data sources and quantify the uncertainty related to these.

### Future invasion risks

Overall (i.e. when considering SRC), we predict that future climate change will reduce invasion risk for *P. grandis* according to both Worldclim and CHELSA, as commonly found for invasive reptiles (Bellard et al. 2013; but see Piquet et al. 2021). Note that in absence of biosecurity measures, a high probability of invasion might persist despite climate effects. On the other hand, invasion risk will increase for *P. laticauda* according to Worldclim but will not change according to CHELSA. For both species, Worldclim-based projections tended to predict higher risks than CHELSA-based projections. The spatial mismatch (overlap) was also greater with Worldclim. The differences between future projections based on CHELSA and Worldclim were of similar, or greater extent to that between the two extreme scenarios (SSP126 and SSP585; Fig. 4). This suggests that the inclusion of multiple climate datasets is of similar importance to that of emission scenarios. Reptiles may shift their phenology in response to environmental change (Kearney et al. 2009), and this has already been observed in *Phelsuma* spp. (Dubos et al. 2020; Baxter-Gilbert et al. 2021). Behavioural response to climate change—and therefore phenological shifts—may interact with geographic response (Kearney et al. 2010). Further research is needed to fully understand the response of invasive reptiles to climate change and improve proactive actions.

### Concluding remarks

The source of climate data was not accounted for in SDM studies until recently (Baker et al. 2016; Morales-Barbero and Vega-Álvarez 2019; Datta et al. 2020; Ocon 2020; Dubos et al. 2022b, a; Stewart et al. 2022). To the best of our knowledge, this study is the first to account for multiple sources of climate data in invasion risk assessments. We highlighted spatial differences in the quantification of environmental suitability, potentially leading to the omission of at-risk regions. Further studies should assess the sensitivity of invasion risks to climate data at broader taxonomic scales, and across different landscapes (especially smaller oceanic islands vs. continents).

The economic cost of IAS is low when detected early, but rises rapidly if not detected because of the damage caused and increased management efforts (Renault et al. 2021). Reptiles represent the second worst invasive vertebrate class in terms of annual economic cost worldwide (Diagne et al. 2021). Therefore, it seems largely economically viable to promote efficient biosecurity measures in order to ensure early detections (Cuthbert et al. 2022) and develop public awareness to reduce intentional release (Perry and Farmer 2011). Given the ecological and economic stakes, surveillance programmes should be considered in areas identified as at high risk of invasion based on single climate data. However, surveillance efforts should be prioritised where high invasion risks are identified with high certainty, i.e., based on predictions accounting for multiple climate data sources.

## Acknowledgements

Portuguese National Funds through FCT (Fundação para a Ciência e a Tecnologia) support the research contract to Angelica Crottini [2020.00823.CEECIND/CP1601/CT0003] and Nicolas Dubos [ICETA_2021_26]. We thank Gregory Deso for useful exchanges and comments on the manuscript.

## Data availability

All biological data formatted and analysed during this study are included in this published article in the supporting online material as RDS objects.

## Declaration

### Author contribution

*All authors contributed to the study conception and design. Material preparation and analyses were performed by Nicolas Dubos. The first draft of the manuscript was written by Nicolas Dubos and all authors commented on previous versions of the manuscript. All authors read and approved the final manuscript*

