## Supporting information for "Choice of climate data influences current and future global invasion risks for two *Phelsuma* geckos"

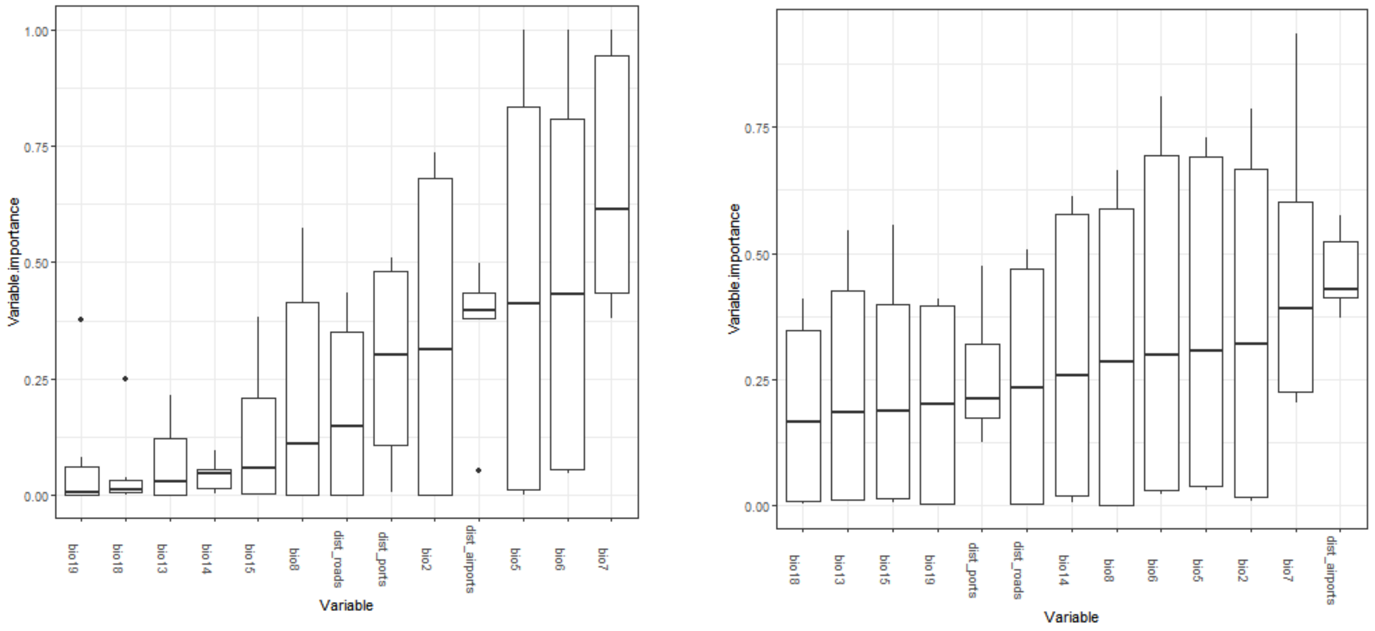


Fig S1. Variable importance for *Phelsuma grandis* based on CHELSA climate data (left: uncorrected models; right: sample bias-corrected models).

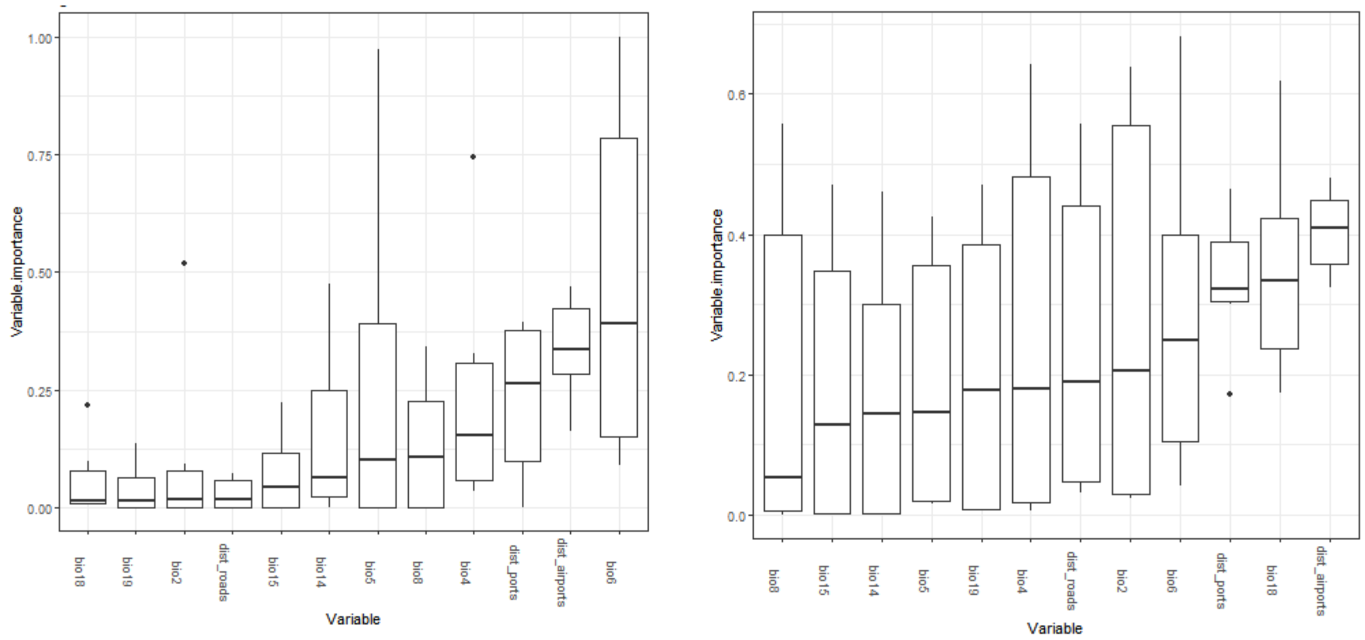


Fig S2. Variable importance for *Phelsuma grandis* based on Worldclim climate data (left: uncorrected models; right: sample bias-corrected models).

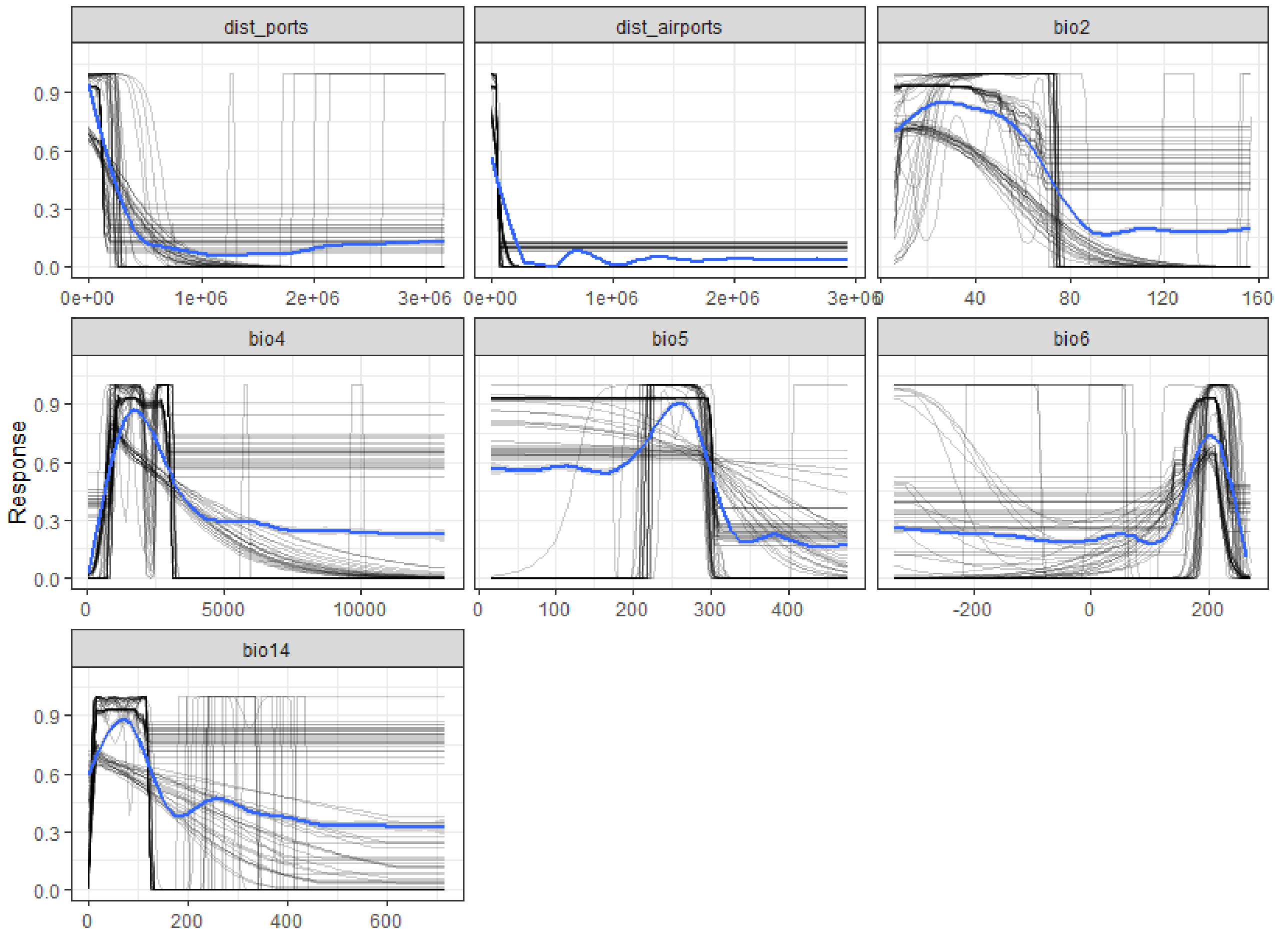


Fig. S3 Response curves for *Phelsuma grandis* distribution models, bias-corrected and calibrated with CHELSA climate data. Black lines are individual response curves for each model iteration, blue lines are smoothed responses.

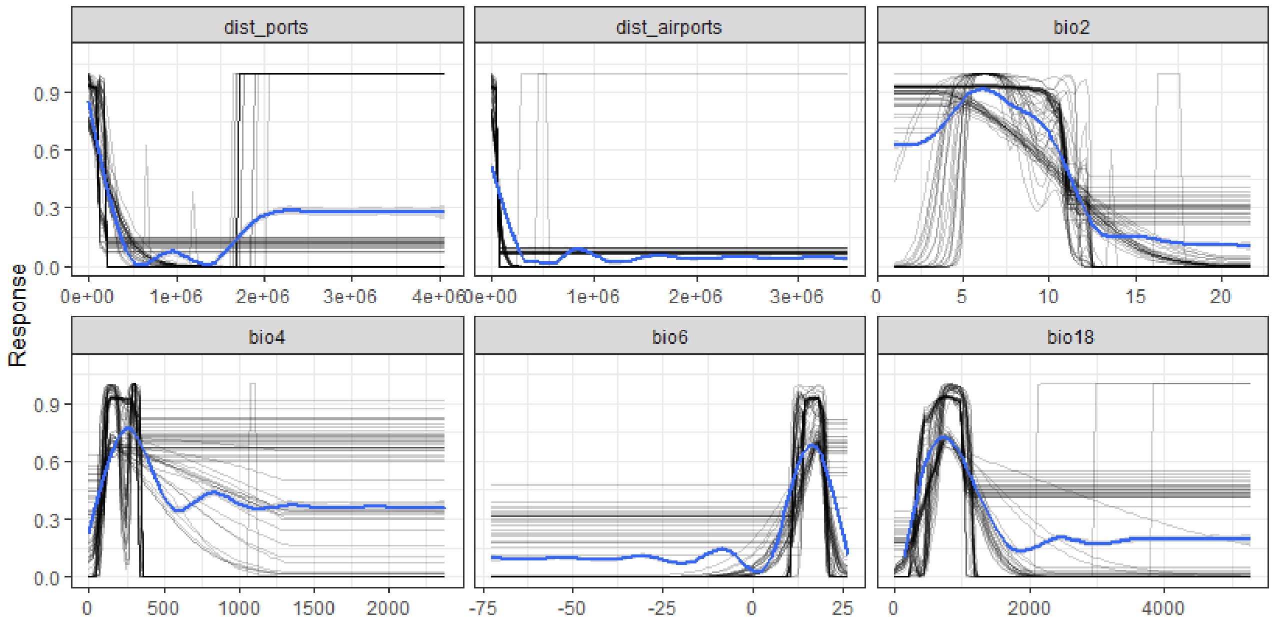


Fig. S4 Response curves for *Phelsuma grandis* distribution models, bias-corrected and calibrated with Worldclim climate data. Black lines are individual response curves for each model iteration, blue lines are smoothed responses.

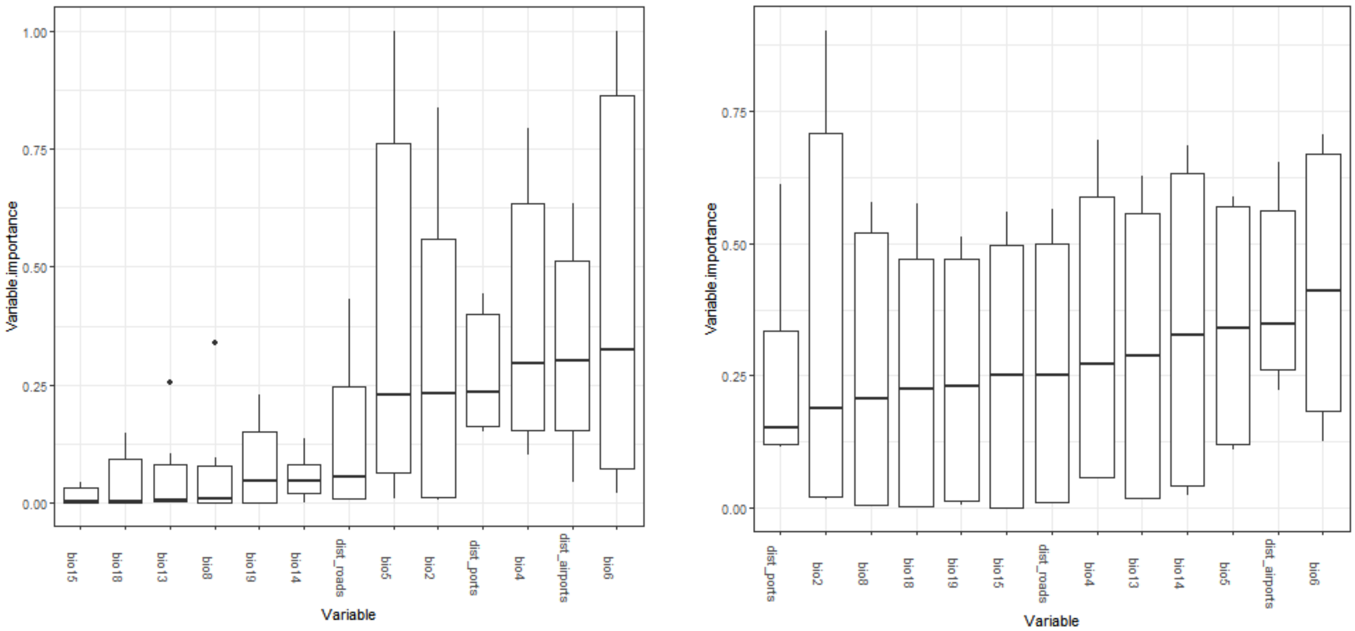


Fig S5. Variable importance for *Phelsuma laticauda* based on CHELSA climate data (left: uncorrected models; right: sample bias-corrected models).

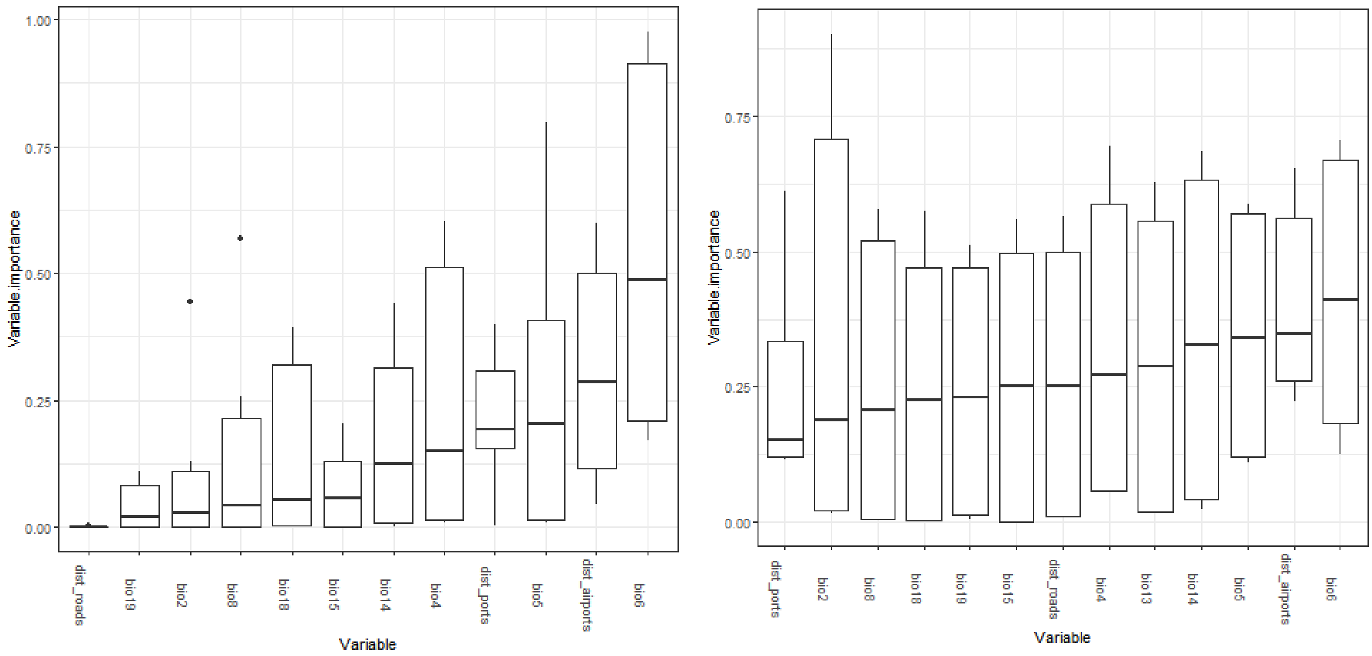


Fig S6. Variable importance for *Phelsuma laticauda* based on Worldclim climate data (left: uncorrected models; right: sample bias-corrected models).

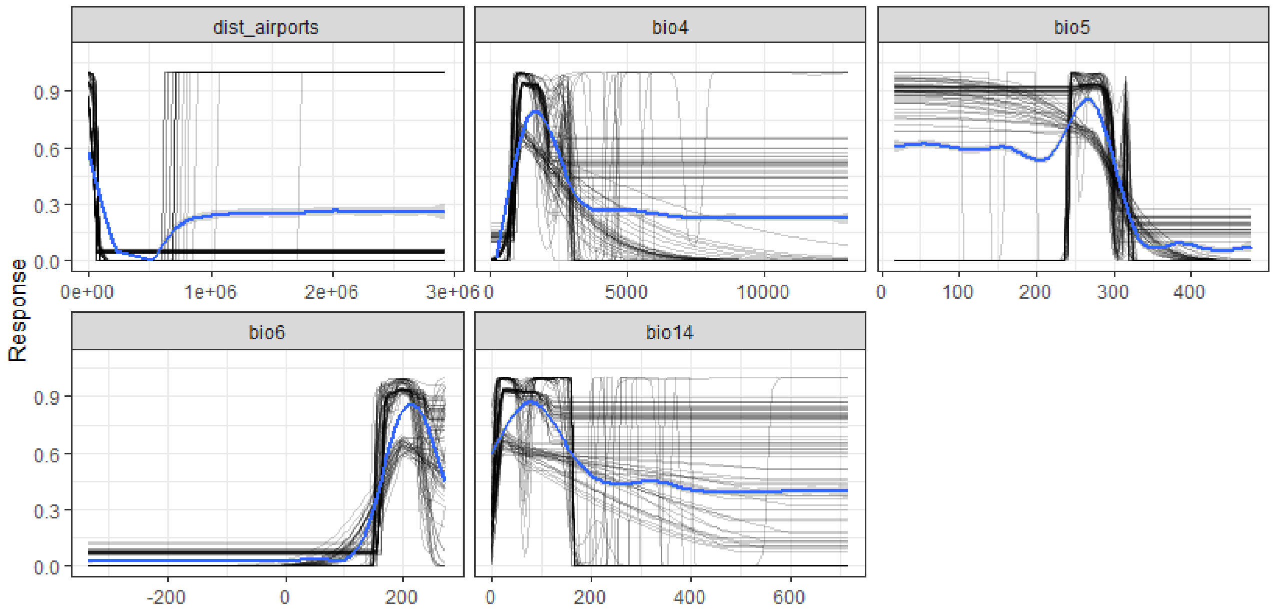


Fig. S7 Response curves for *Phelsuma laticauda* distribution models, bias-corrected and calibrated with CHELSA climate data. Black lines are individual response curves for each model iteration, blue lines are smoothed responses.

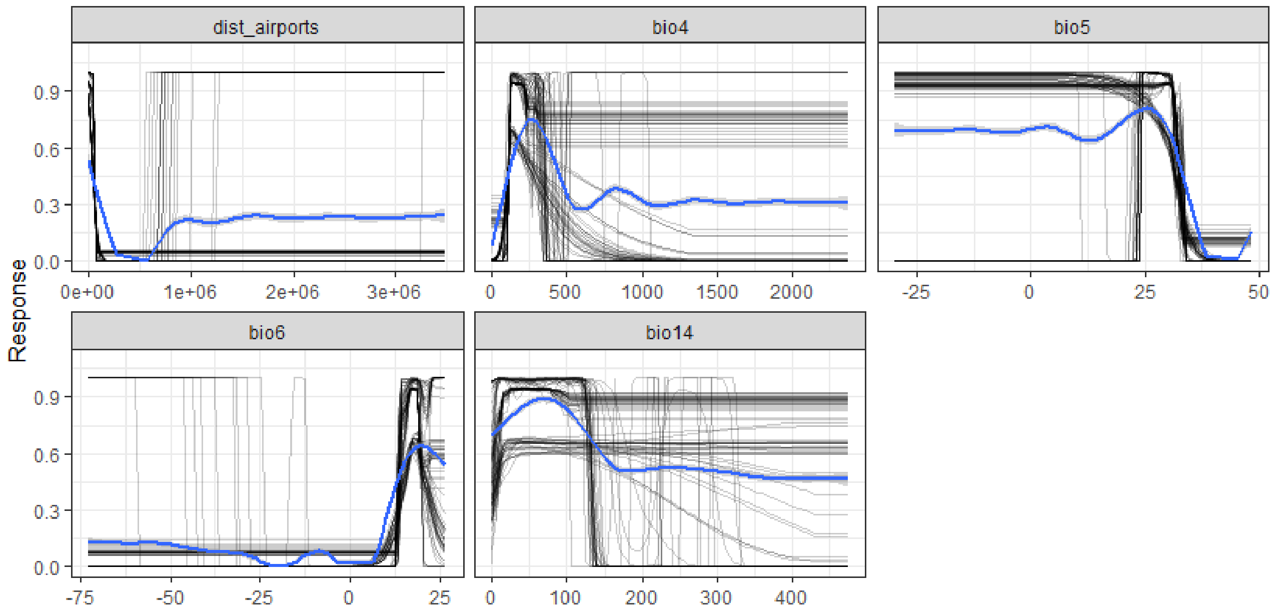


Fig. S8 Response curves for *Phelsuma laticauda* distribution models, bias-corrected and calibrated with Worldclim climate data. Black lines are individual response curves for each model iteration, blue lines are smoothed responses.

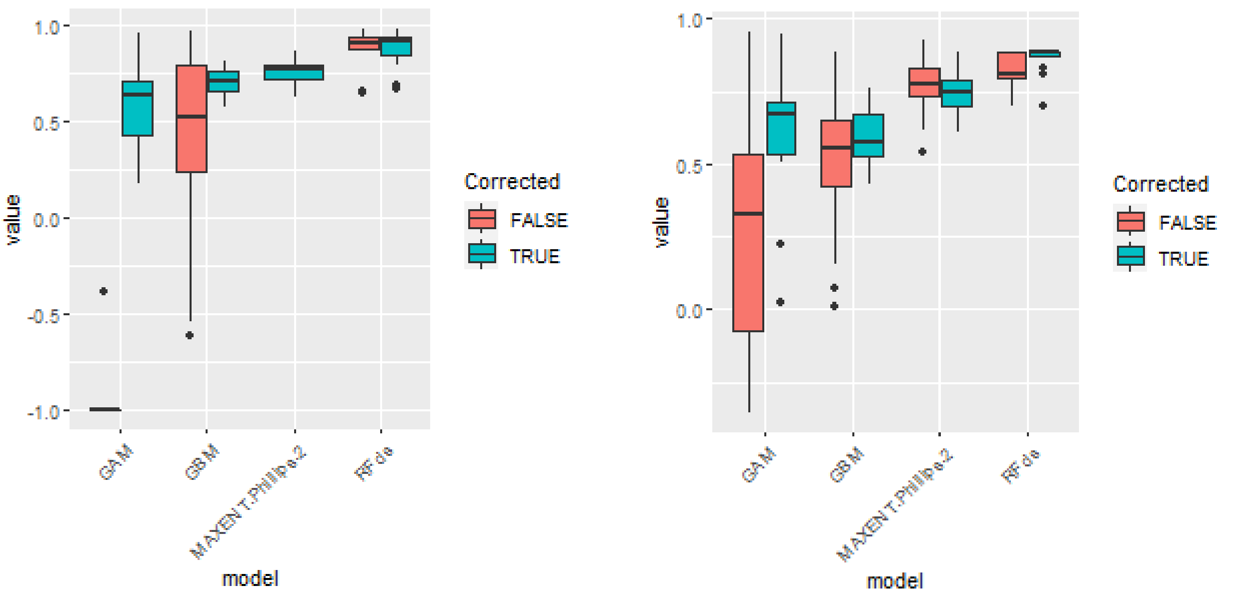


Fig. S9 Model performance for *P. grandis* (Boyce index) for four modelling techniques individually, for two modalities of sample bias correction (red: uncorrected; blue: corrected) and two climate data (left: CHELSA; right: Worldclim). Boxes represent 25-75% quantiles and the bar represent the median.

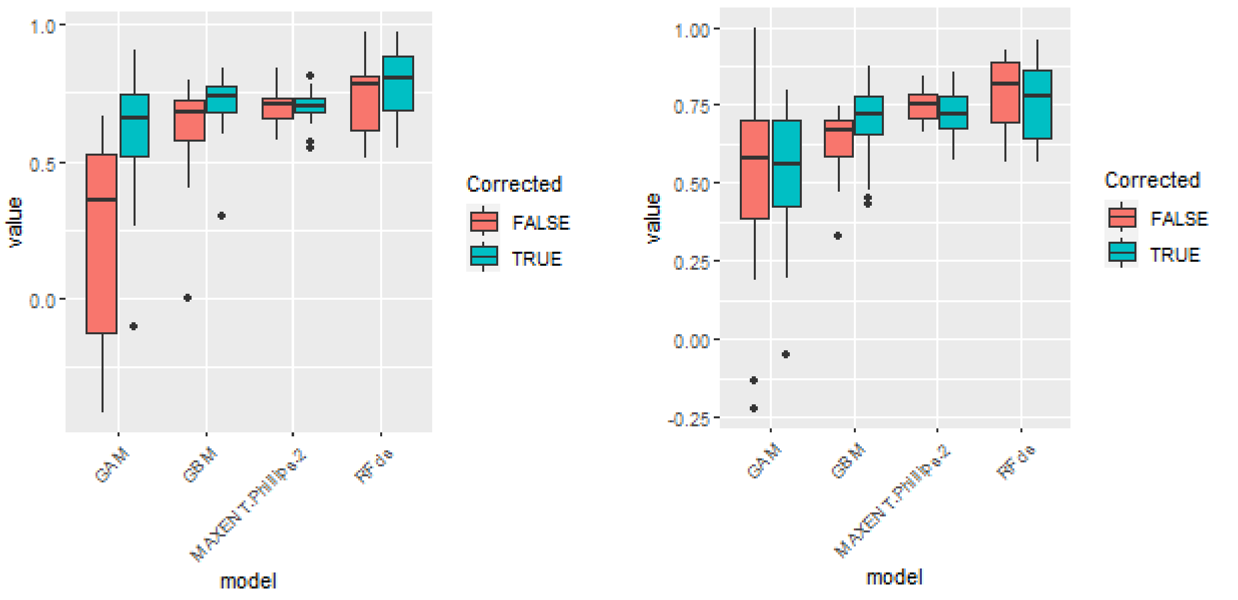


Fig. S10 Model performance for *P. laticauda* (Boyce index) for four modelling techniques individually, for two modalities of sample bias correction (red: uncorrected; blue: corrected) and two climate data (left: CHELSA; right: Worldclim). Boxes represent 25-75% quantiles and the bar represent the median.

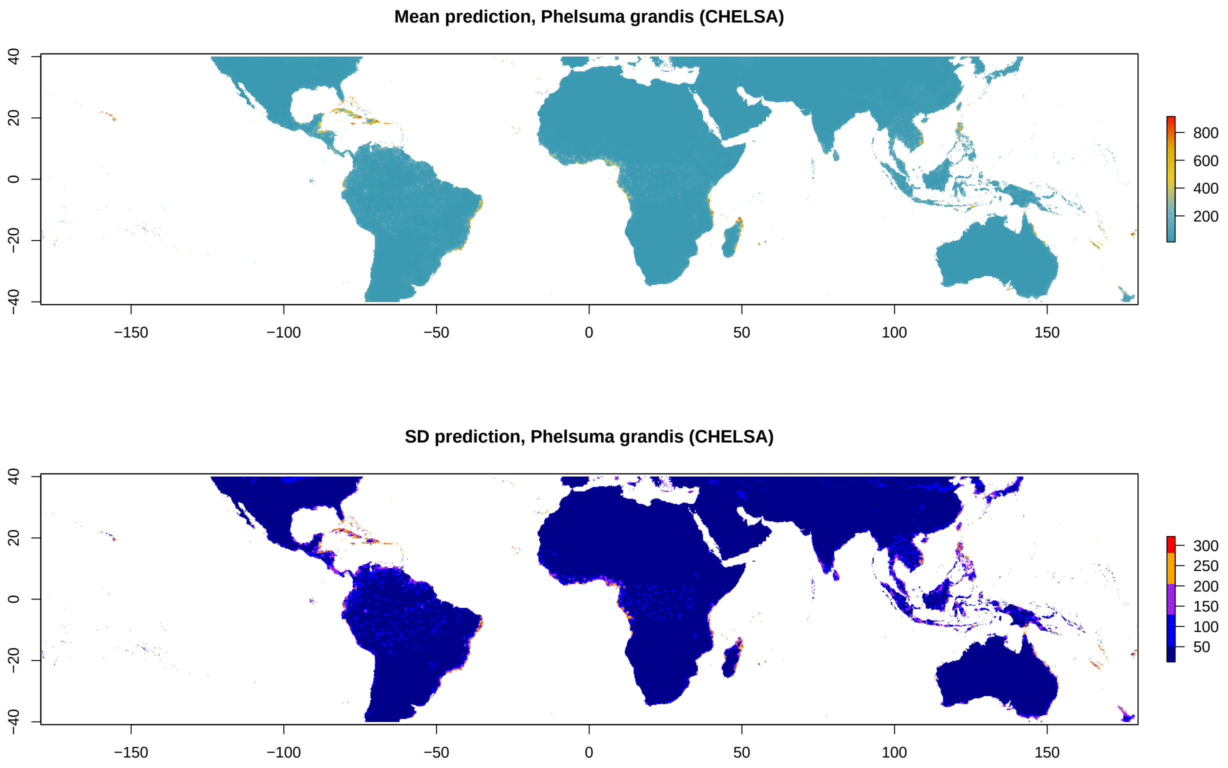


Fig. S11. Mean (top) and standard deviation (bottom) prediction of current invasion risk based on CHELSA climate data across high performing models for *Phelsuma grandis*.

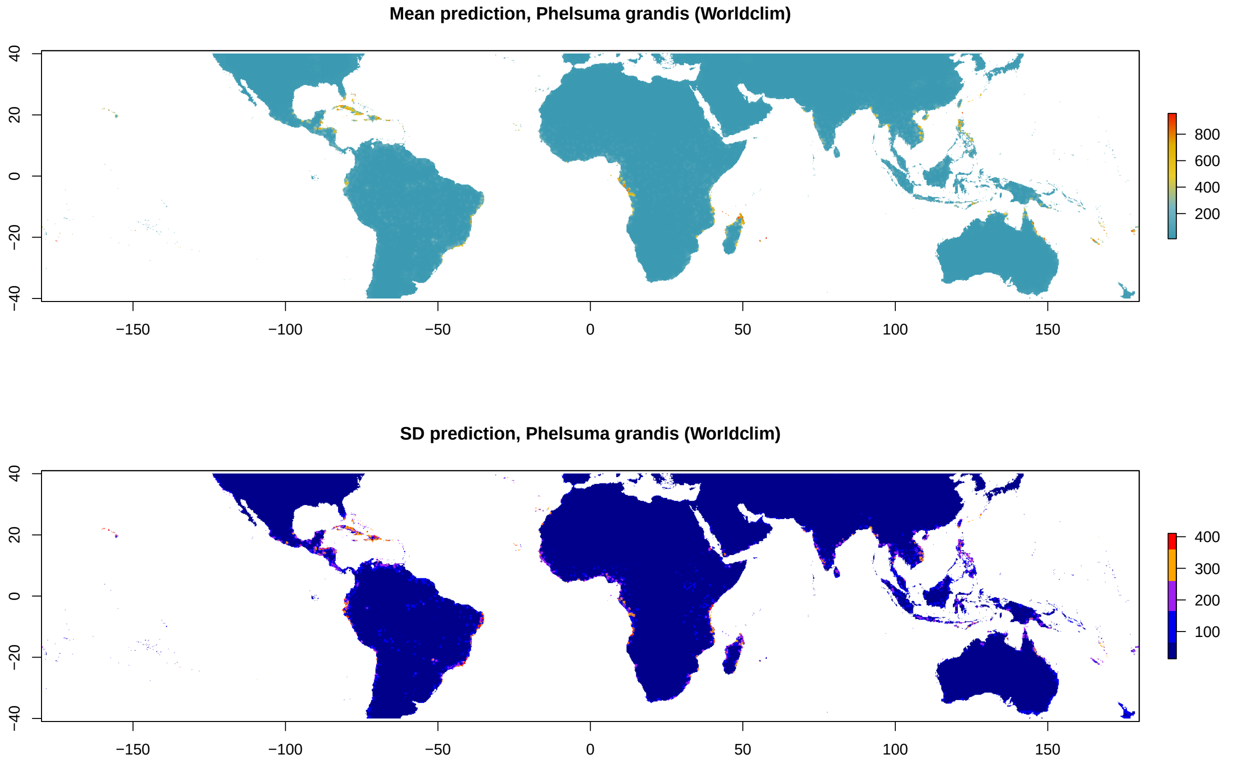


Fig. S12. Mean (top) and standard deviation (bottom) prediction of current invasion risk based on Worldclim climate data across high performing models for *Phelsuma grandis*.

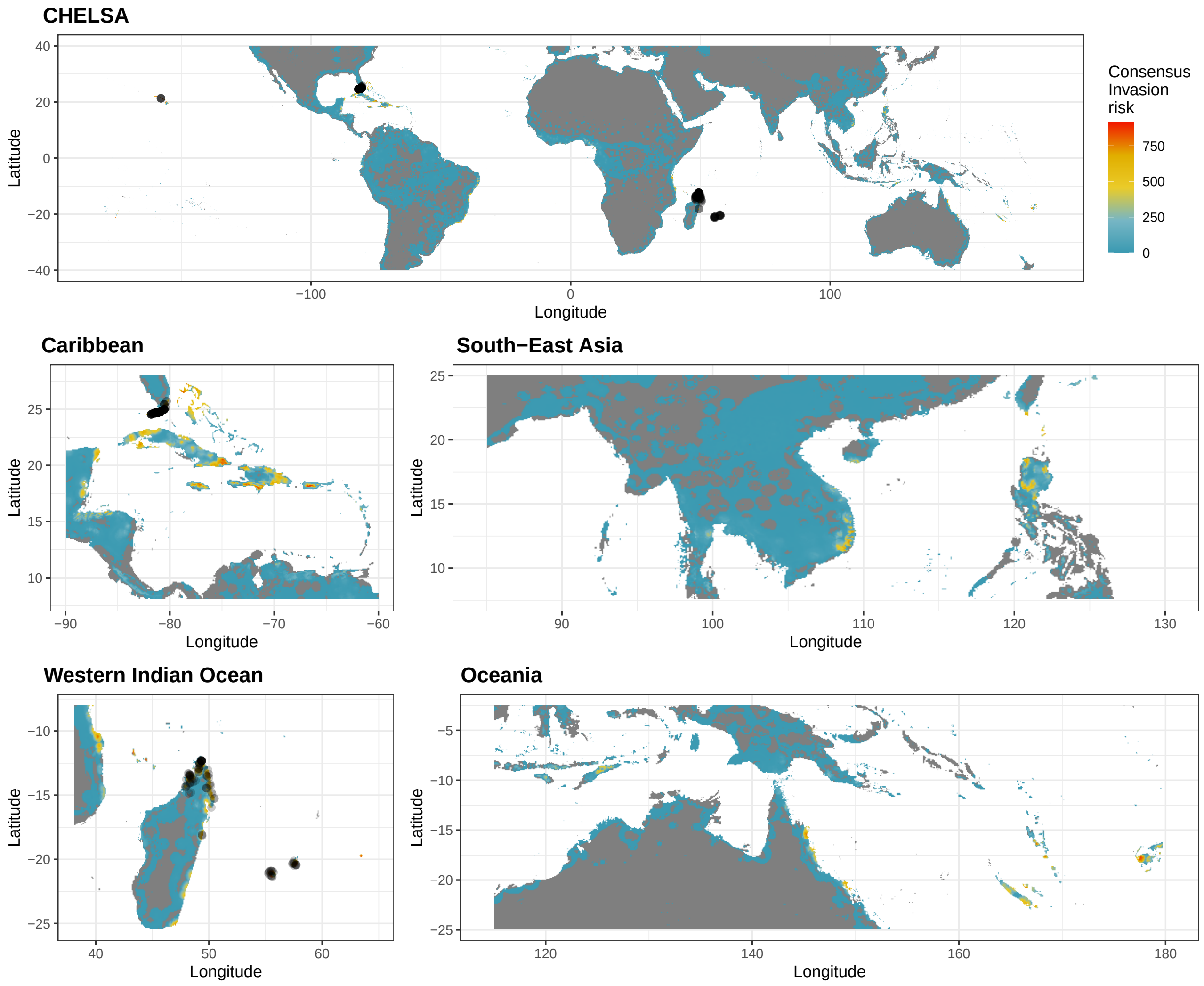


Fig. S13. Consensus invasion risk (mean prediction – standard deviation across model replicates) for *Phelsuma grandis* based on CHELSA climate data only.

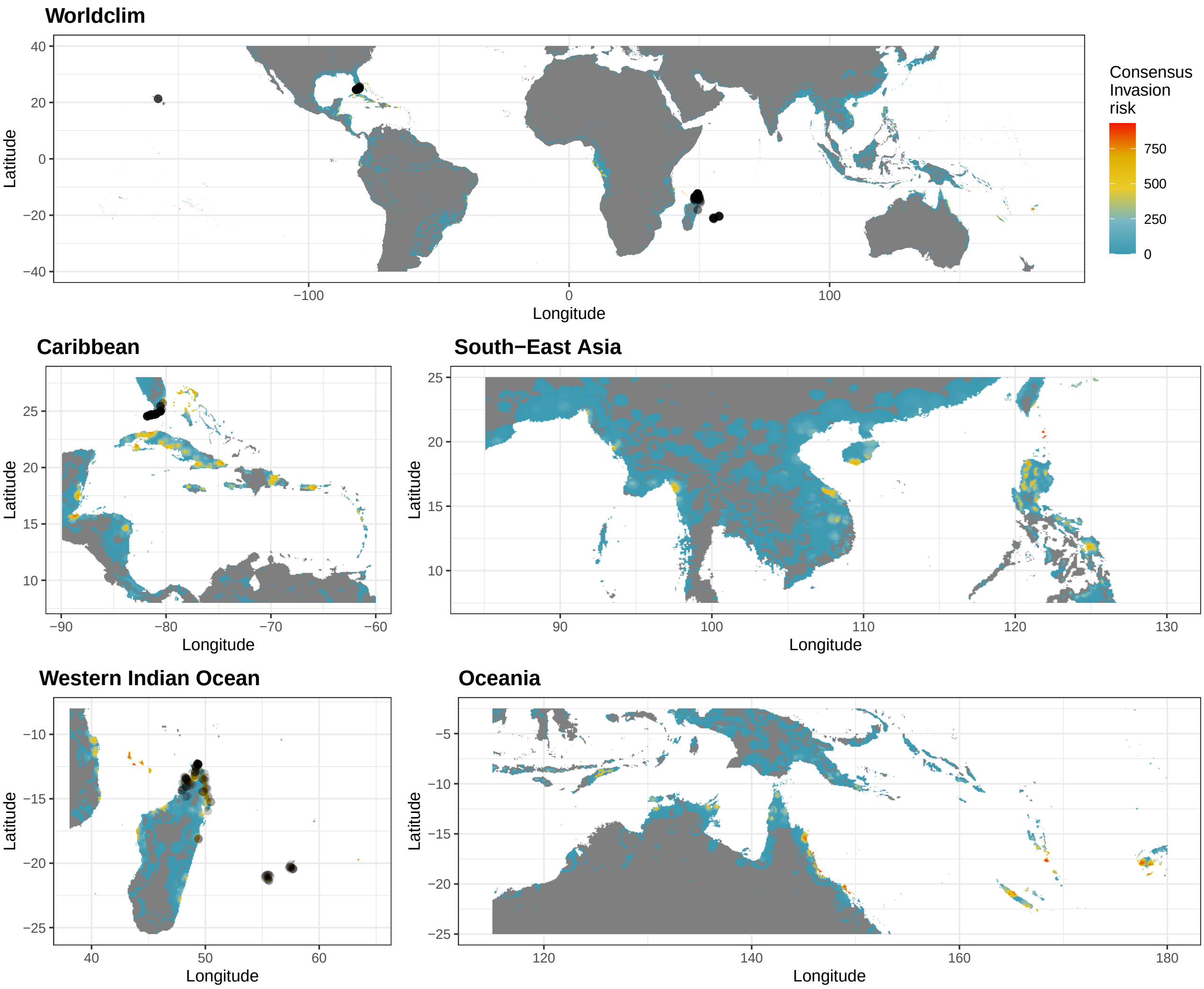


Fig. S14 Consensus invasion risk (mean – standard deviation across model replicates) for *Phelsuma grandis* based on Worldclim climate data only.

Fig. S15 Mean (top) and standard deviation (bottom) prediction of invasion risk based on CHELSA climate data across high performing models for *Phelsuma laticauda*.

Fig. S16 Mean (top) and standard deviation (bottom) prediction of invasion risk based on Worldclim climate data across high performing models for *Phelsuma laticauda*.

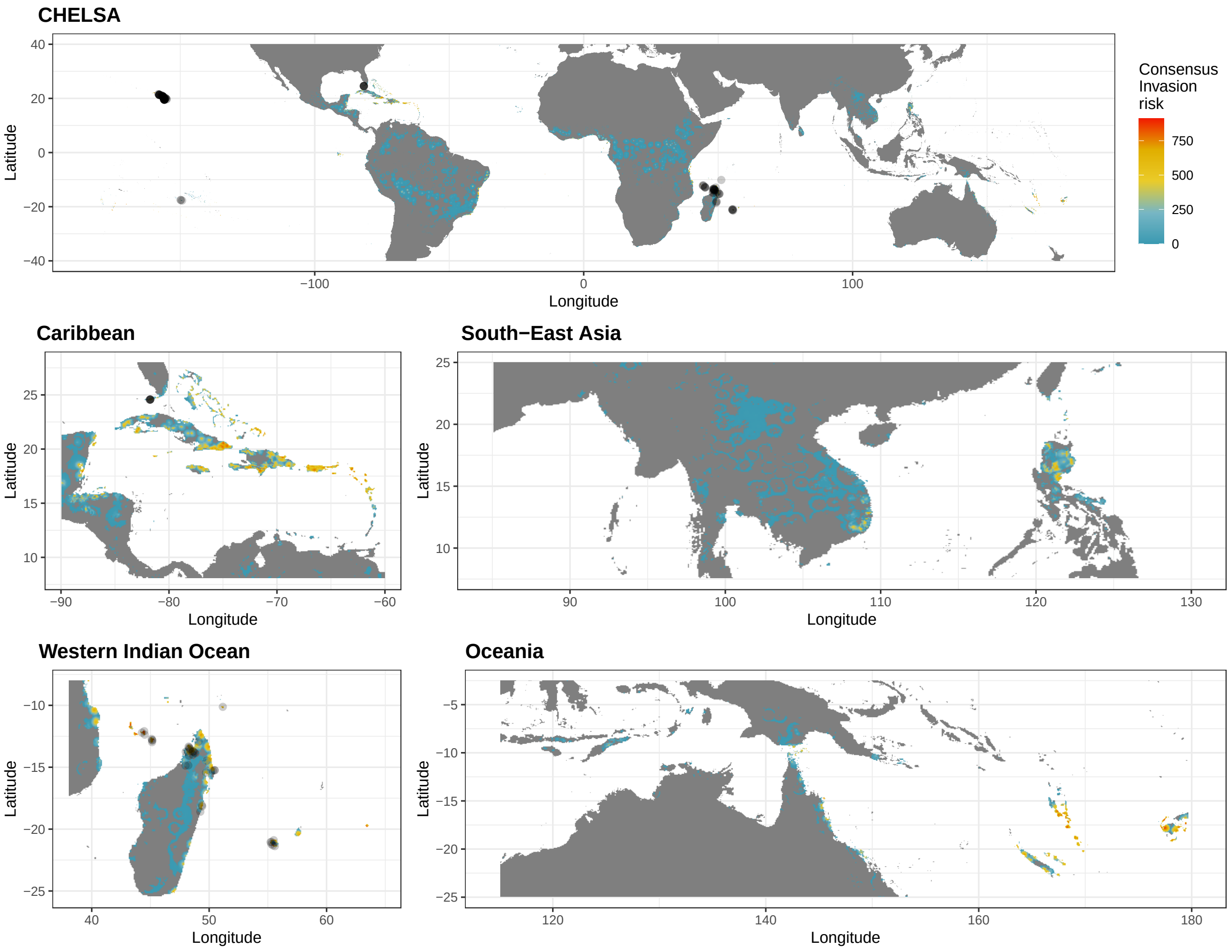


Fig. S17 Consensus invasion risk (mean – standard deviation across model replicates) for *Phelsuma laticauda* based on CHELSA climate data only.

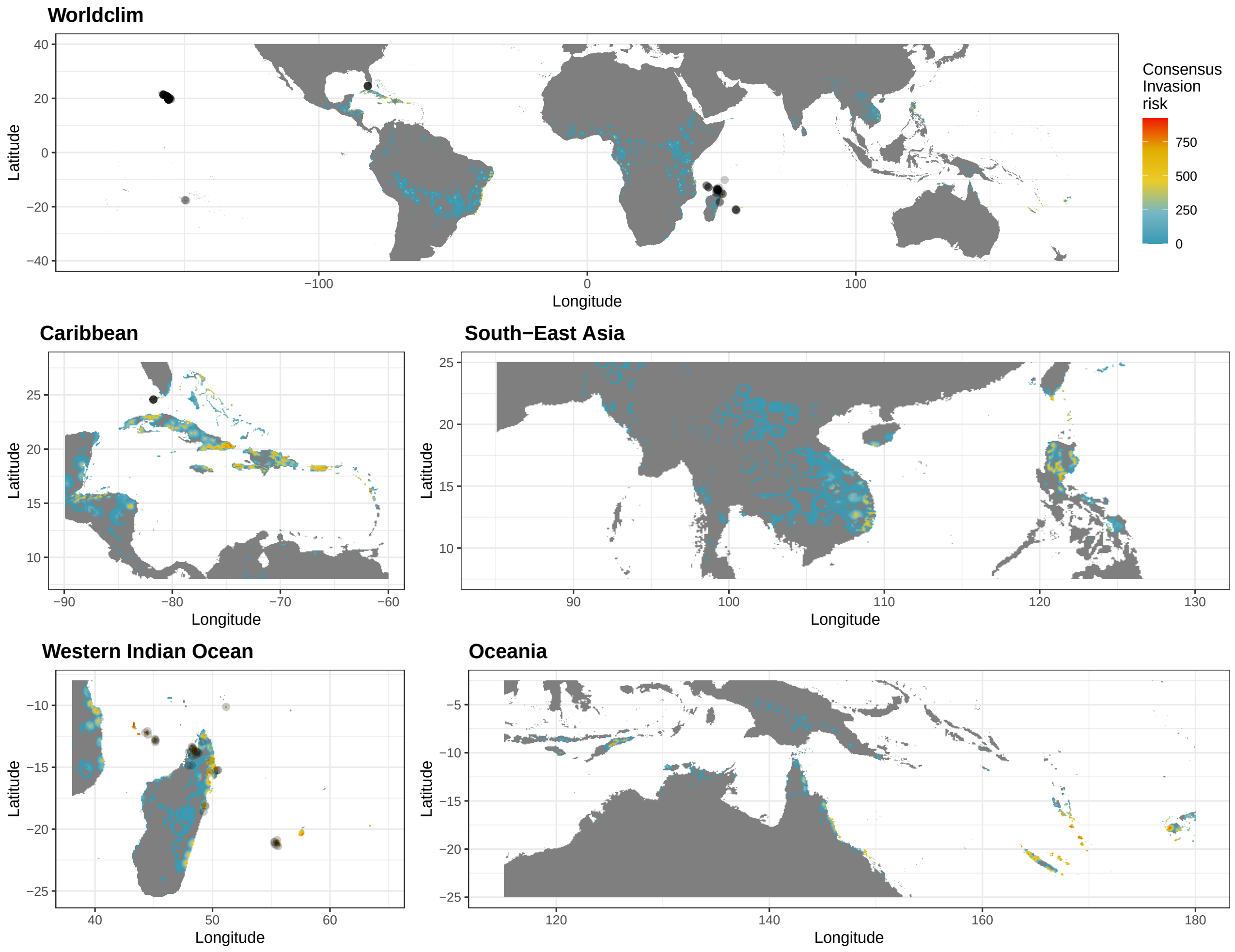


Fig. S18 Consensus invasion risk (mean – standard deviation across model replicates) for *Phelsuma laticauda* based on Worldclim climate data only.

**Future projections**

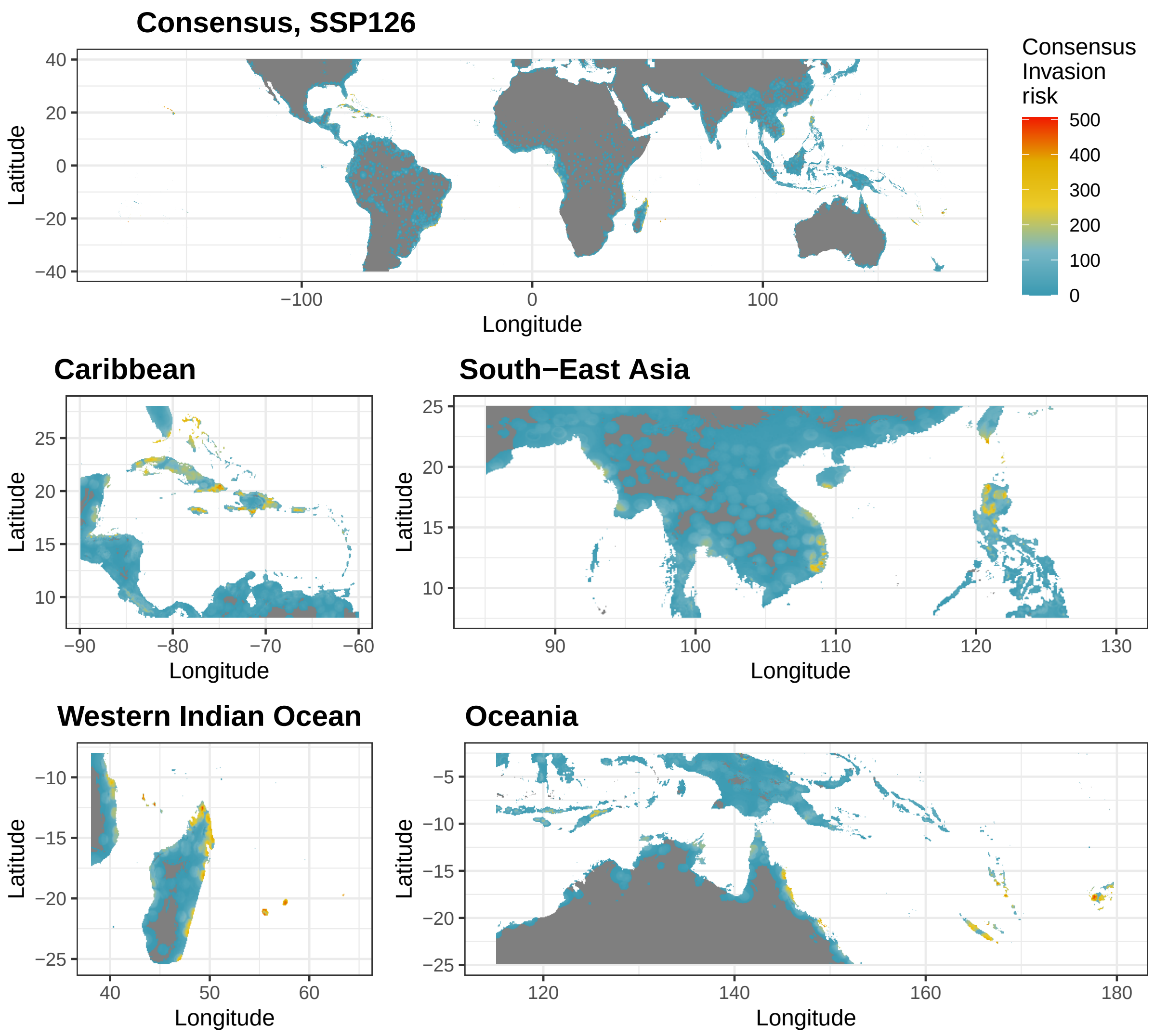


Fig S19 Consensus future invasion risk (2070) for *Phelsuma grandis* according to the SSP126 scenario. Projections were obtained from the mean projection between two climate data sources (CHELSA and Worldclim) and three GCMs, and were penalised by their uncertainty (mean – standard deviation).

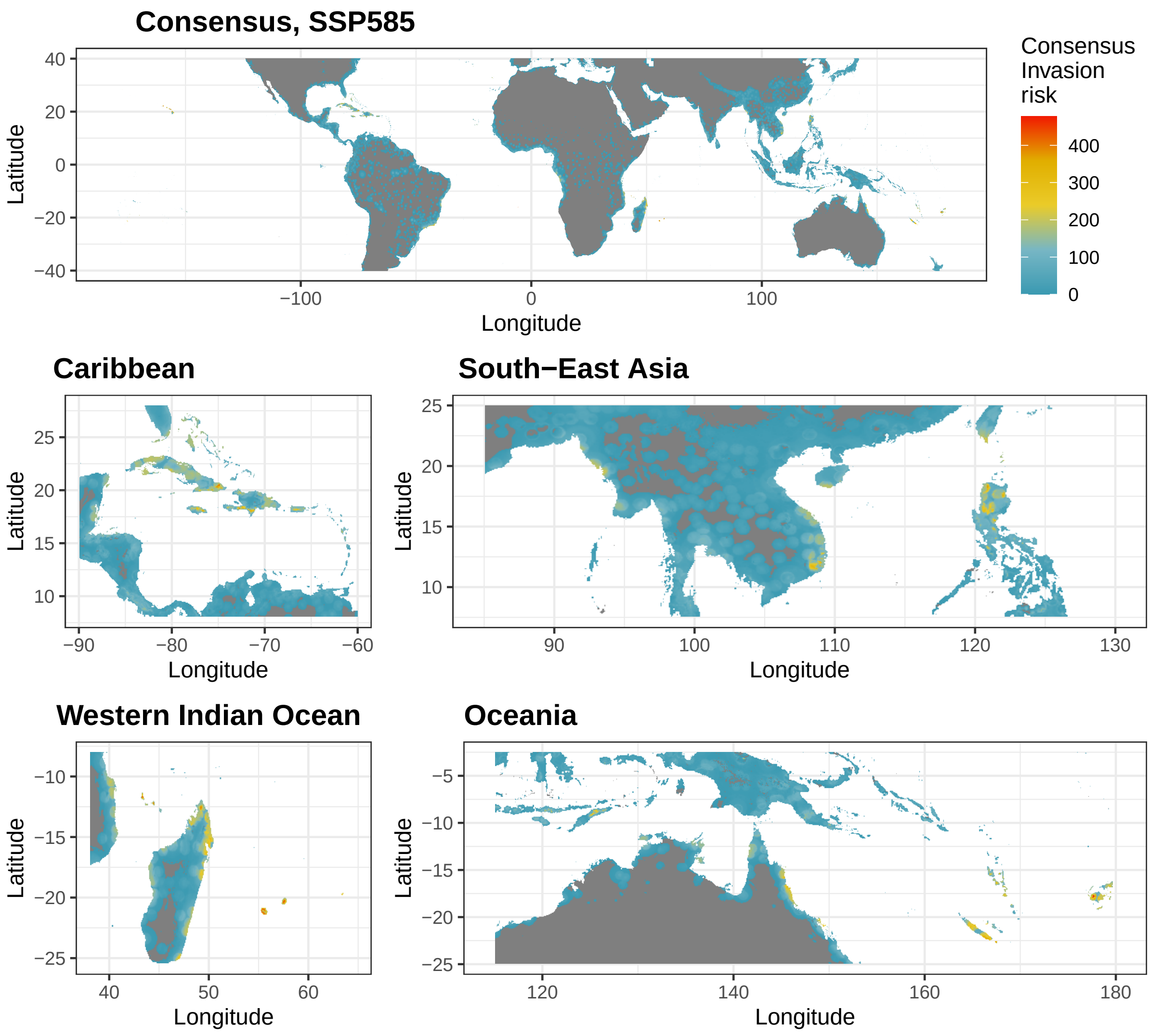


Fig S20 Consensus future invasion risk (2070) for *Phelsuma grandis* according to the SSP585 scenario. Projections were obtained from the mean projection between two climate data sources (CHELSA and Worldclim) and three GCMs, and were penalised by their uncertainty (mean – standard deviation).

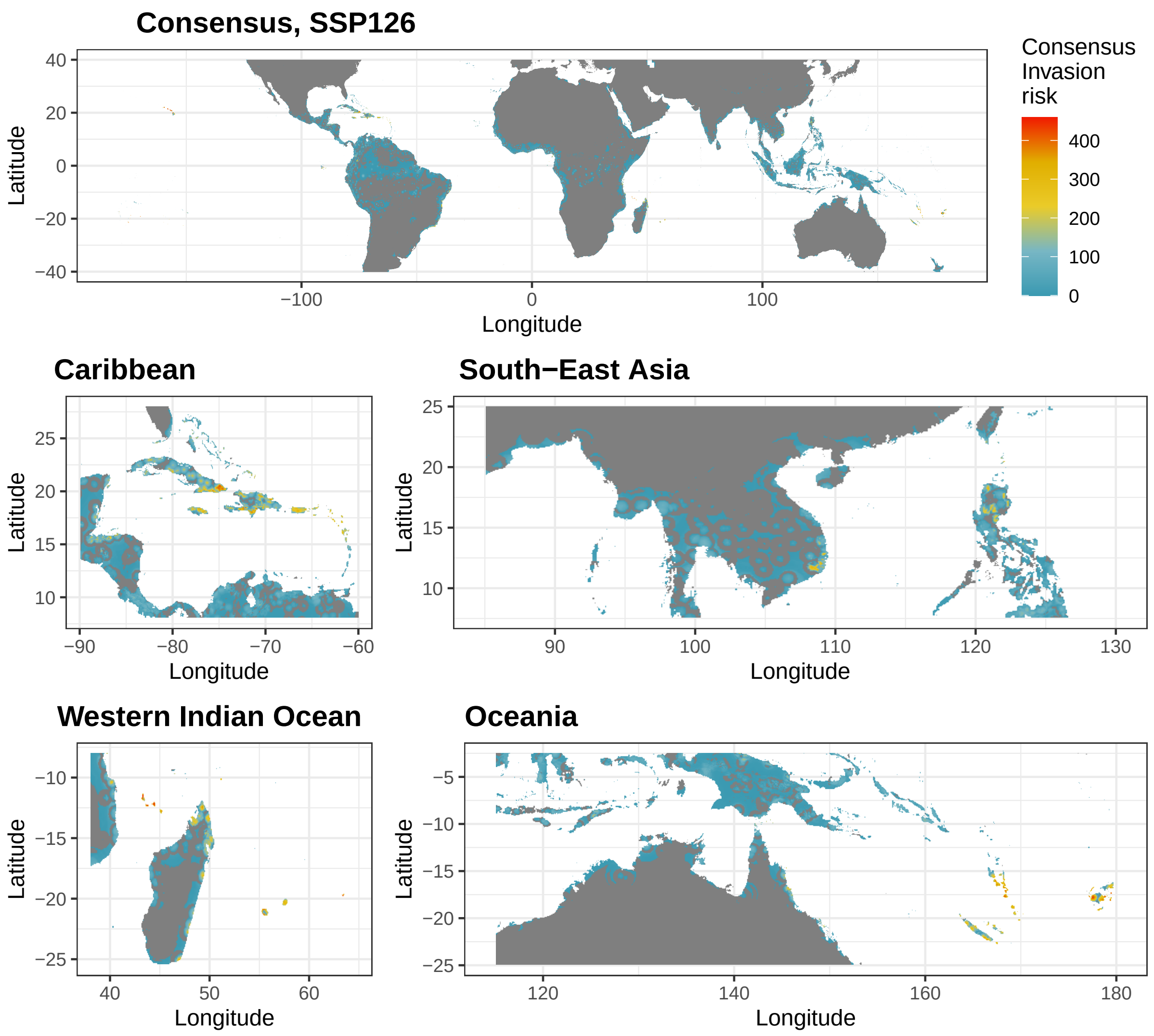


Fig S21 Consensus future invasion risk (2070) for *Phelsuma laticauda* according to the SSP126 scenario. Projections were obtained from the mean projection between two climate data sources (CHELSA and Worldclim) and three GCMs, and were penalised by their uncertainty (mean – standard deviation).

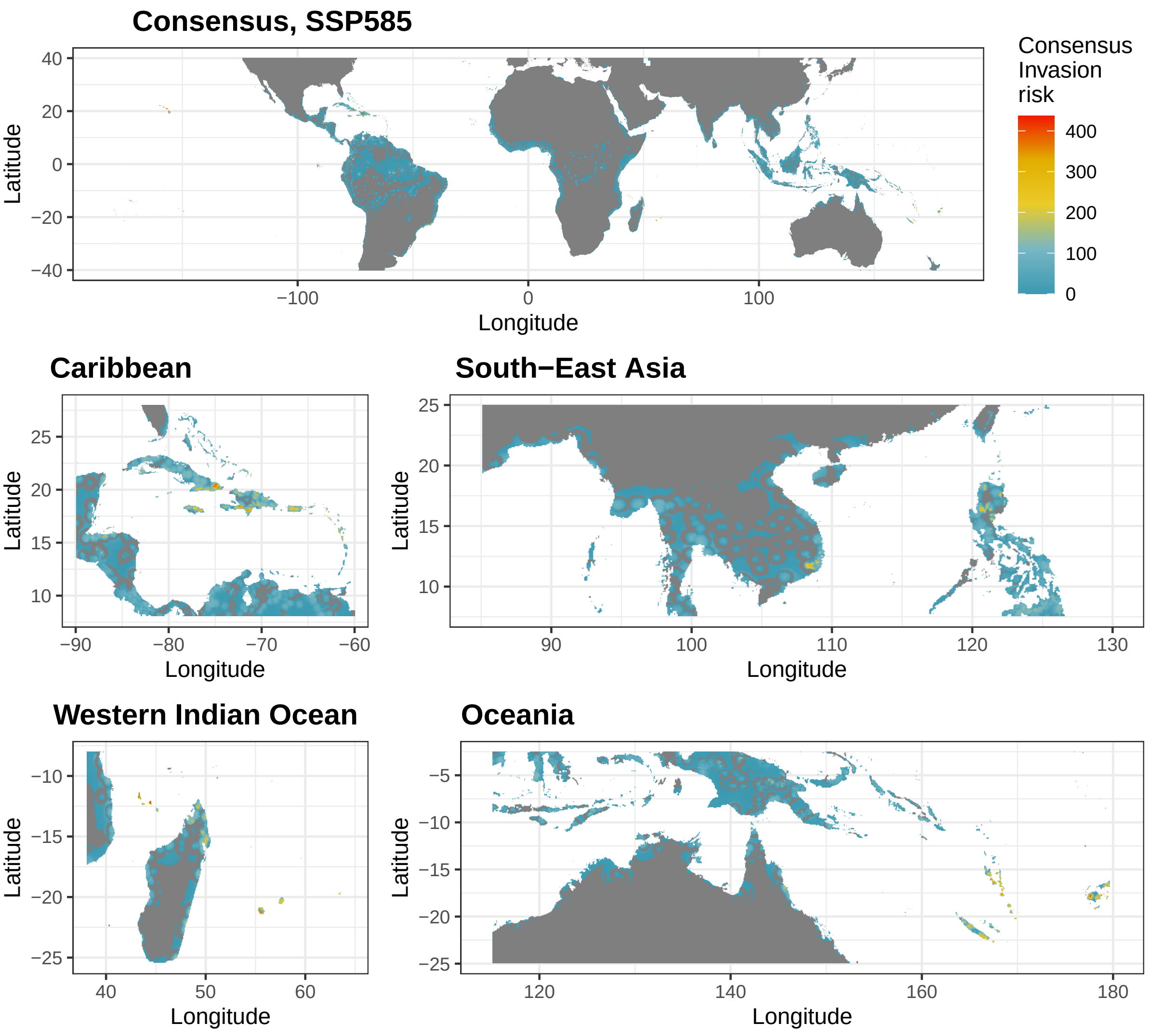


Fig S22 Consensus future invasion risk (2070) for *Phelsuma grandis* according to the SSP585 scenario. Projections were obtained from the mean projection between two climate data sources (CHELSA and Worldclim) and three GCMs, and were penalised by their uncertainty (mean – standard deviation).

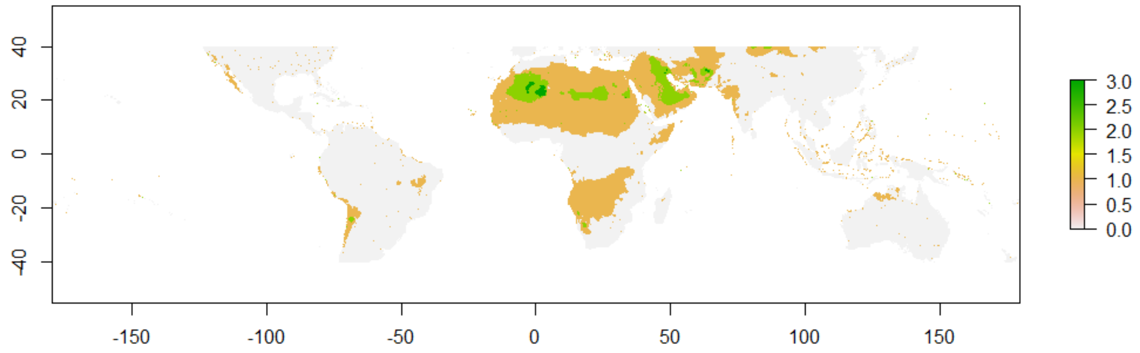


Fig. S23 Example of clamping mask for HadGem2-AO SSP585 from CHELSA for the year 2070. The scale indicates the number of predictors for which novel conditions are met.

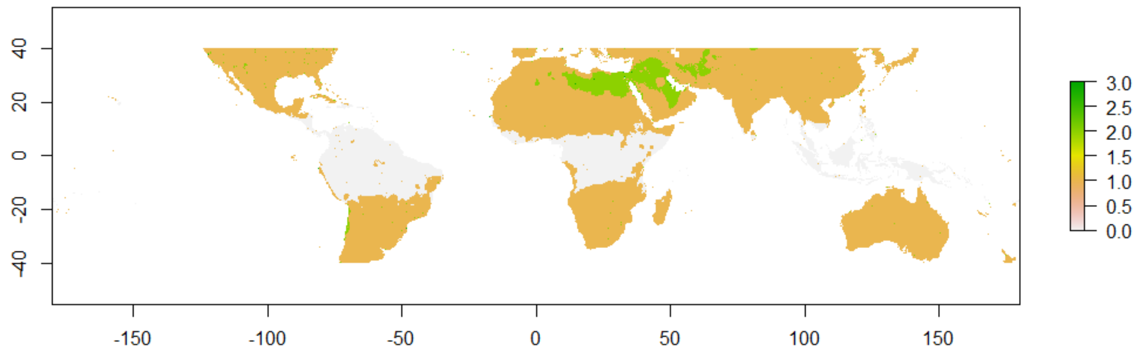


Fig. S24 Example of clamping mask for HadGem2-AO SSP585 from Worldclim for the year 2070. The scale indicates the number of predictors for which novel conditions are met.

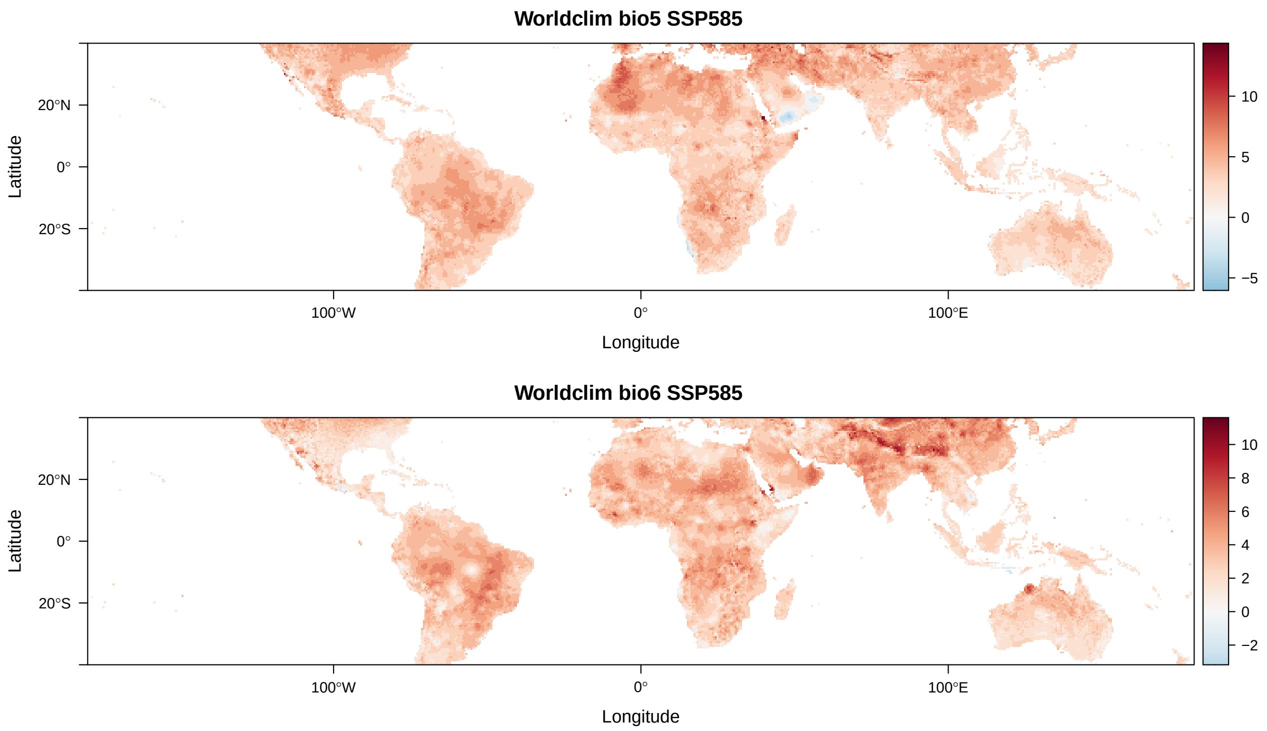


Fig. S25 Differences between current and 2070 conditions for HadGem2-AO SSP585 from Worldclim for bio5 and bio6.
